# Fusion-driven post-transcriptional network orchestrates ferroptosis resistance, dormancy, and immune remodeling in PRCC-TFE3 renal cell carcinoma

**DOI:** 10.64898/2026.02.12.692412

**Authors:** Divya Mishra, Shivangi Agrawal, Divya Malik, Ekta Pathak, Rajeev Mishra

## Abstract

TFE3-rearranged papillary renal cell carcinoma (pRCC) remains poorly understood due to its rarity and complex biology. By integrating matched mRNA and miRNA sequencing from the TCGA-KIRP cohort, this study reveals a fusion-specific post-transcriptional program orchestrating ferroptosis resistance, metabolic adaptation, adhesion loss, immune remodeling, and dormancy-associated signaling. Fusion-positive tumors exhibited coordinated upregulation of antioxidant and mTORC1 components (SOD2, GPX4, G6PD, NQO1, FNIP2, RRAGC), reinforced by loss of miRNA-mediated repression. In parallel, elevated miRNAs (miR-185, miR-148a, miR-130b, miR-342) suppressed MAPK, Wnt, and tight-junction genes, generating a low-proliferative, adhesion-deficient state. AGO2-compatible miRNA-mRNA structural modeling validated key duplexes, linking miRNA regulatory specificity to fusion-driven transcriptional outputs. Immune profiling revealed reduced CD3⁺ T cells but enrichment of CD8⁺ and NK cells, supporting a remodeled yet persistence-permissive microenvironment. Finally, connectivity mapping analysis identified candidate leads targeting the MAPK–Wnt axis. These findings define a multilayered RNA regulatory architecture unique to PRCC-TFE3 tumors.

## Introduction

Renal cell carcinoma (RCC) represents a heterogeneous group of epithelial malignancies, with clear cell (∼70%), papillary (∼15–20%), and chromophobe (∼5–10%) being the most common subtypes [1, 2]. Within papillary RCC (pRCC), a rare yet clinically aggressive subset, translocation RCC (tRCC), is characterized by TFE3 gene rearrangements at Xp11.2 that produce chimeric transcription factors (e.g., PRCC-TFE3, NONO-TFE3, ASPSCR1-TFE3) and result in aberrant transcriptional activity [3–6]. Among these, PRCC-TFE3 is one of the most frequently observed events in pRCC and is associated with distinctive histopathology and adverse clinical behavior [7, 8]. Recent studies have expanded our understanding of tRCC molecular biology by implicating TFE3 protein fusions in multiple dysregulated pathways, including mTOR signaling, oxidative phosphorylation, ferroptosis resistance, epithelial–mesenchymal transition (EMT), and immune modulation [9, 10]. In particular, TFE3 fusion proteins activate mitochondrial biogenesis and mitophagy, supporting metabolic adaptation and limiting oxidative stress [7, 9, 11]. However, the mechanisms by which these transcriptional programs are fine-tuned at the post-transcriptional level remain insufficiently defined. Moreover, despite the use of mTOR inhibitors and tyrosine kinase inhibitors (TKIs) in TFE3-rearranged RCC, clinical benefit has been variable [12–14], highlighting the need to identify molecular regulators and biomarkers that distinguish fusion-positive from fusion-negative pRCC.

MicroRNAs (miRNAs) are small non-coding RNAs that downregulate gene expression by binding to partially complementary sequences in the 3′ untranslated region (3′UTR) of target transcripts, thereby promoting translational repression and/or transcript degradation [15–17]. A single miRNA can regulate numerous genes, and conversely, individual genes may be targeted by multiple miRNAs, creating interconnected miRNA–mRNA regulatory networks that coordinate tumor-associated processes. Dysregulated miRNA expression has been associated with cancer-related pathways such as cell proliferation, apoptosis, metabolism, and immune evasion [18–23]. In RCC, specific miRNAs have been shown to modulate ferroptosis, mTOR signaling, and immune infiltration, but these interactions have seldom been assessed in the context of TFE3 fusion status [24–30].

The tumor microenvironment (TME) influences disease progression and therapeutic response through multifaceted crosstalk between tumor cells, immune populations, and stromal elements [31–36]. Understanding how miRNA–mRNA interactions shape both intracellular signaling and the surrounding immune milieu is thus important for defining the molecular heterogeneity of pRCC.

In this study, we analyzed matched mRNA and miRNA sequencing from the TCGA-KIRP cohort to delineate gene regulatory networks that distinguish PRCC-TFE3 fusion-positive from fusion-negative pRCC. This combined analysis identified a fusion-specific miRNA–mRNA architecture connecting ferroptosis resistance, metabolic reprogramming, dormancy-associated transcriptional suppression, and immune remodeling. By applying AGO2-centered structural modelling [37] together with ROC-based biomarker evaluation, we corroborated key post-transcriptional interactions and identified fusion-dependent regulatory susceptibilities. Collectively, these results indicate that PRCC-TFE3 fusion reshapes RNA-level regulation across metabolic, structural, and immune pathways, providing mechanistic insight into the molecular divergence between fusion-positive and fusion-negative tumors and highlighting potential diagnostic markers and therapeutic targets for this aggressive renal cancer subtype.

## Results

### Transcriptomic profiling of PRCC-TFE3 fusion and fusion-negative KIRP:TCGA cohort

RNA-seq and miRNA-seq datasets from the TCGA-KIRP cohort were categorized into fusion-positive and fusion-negative pRCC cases, with normal kidney tissues serving as controls (Figure 1A). Differential gene expression analyses were performed using both DESeq2 and LIMMA methods. Although the two methods produced partially distinct gene lists, their intersection identified 2,117 DEGs (1,298 upregulated and 819 downregulated) for the fusion-positive group and 3,348 DEGs (1,399 upregulated and 1,949 downregulated) for the fusion-negative group (Figure 1B and C). The intersection of DESeq2 and LIMMA outputs yielded a consensus DEG set for downstream analysis (Figure 1C). Comparison of the method-specific DEG subsets showed 880 upregulated and 301 downregulated genes specific to fusion-positive KIRP, and 981 upregulated and 1,431 downregulated genes specific to fusion-negative KIRP (Figure 1C). These transcriptional patterns indicate marked molecular divergence between PRCC-TFE3 fusion-positive and fusion-negative papillary renal carcinomas and provide a basis for subsequent network and pathway enrichment analyses.

**Figure 1.**
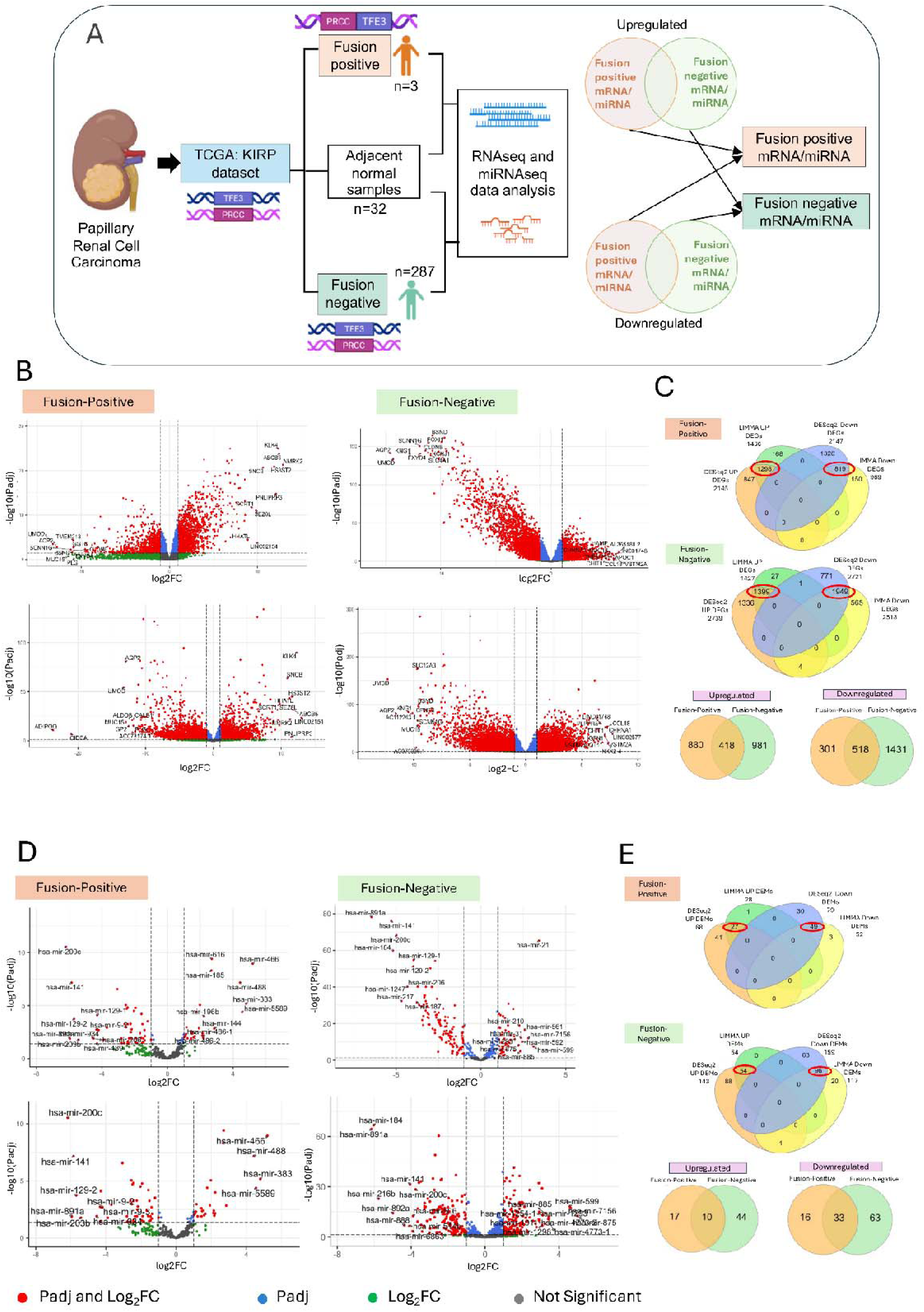
Differential expression analysis of mRNAs and miRNAs in PRCC-TFE3 fusion-positive and fusion-negative pRCC. **(A)** Schematic overview of the analytical workflow. RNA-seq and miRNA-seq datasets from the TCGA-KIRP cohort were categorized into fusion-positive and fusion-negative pRCC cases, with normal kidney tissues serving as controls. DEGs and DEMs were identified using LIMMA and DESeq2, and intersecting and exclusive sets for each fusion group were derived using Venn analysis. Created with Biorender.com. **(B**) Volcano plots showing DEGs obtained from fusion-positive (left) and fusion-negative (right) datasets using both LIMMA (upper panel) and DESeq2 (lower panel). Significantly upregulated and downregulated genes are highlighted in red; blue and green points indicate genes significant only by adjusted p-value (padj) or log_₂_ fold change (log_₂_FC), respectively; grey points denote non-significant genes.(**C).**Venn diagrams depicting overlap of upregulated and downregulated DEGs identified by LIMMA and DESeq2 for each fusion group. Intersections represent high-confidence DEGs used for downstream pathway enrichment; the lower panels show fusion-group–specific exclusive upregulated and downregulated gene sets in fusion-positive and fusion-negative pRCC.**(D)**.Volcano plots of differentially expressed miRNAs (DEMs) in fusion-positive and fusion-negative pRCC. **(E)** Venn diagrams summarizing overlap and exclusivity of DEMs detected by LIMMA and DESeq2. The upper panels show intersecting upregulated and downregulated miRNAs in each fusion group, while the lower panels illustrate fusion-group–specific miRNAs identified in fusion-positive and fusion-negative pRCC cohorts.

### Differential expression of miRNAs in the TCGA-KIRP cohort

Differential miRNA expression analysis was performed for PRCC-TFE3 fusion-positive pRCC and fusion-negative pRCC relative to normal samples using both LIMMA and DESeq2 methods. The resulting DEM profiles for both datasets are shown as volcano plots (Figure 1D). Intersecting the DESeq2 and LIMMA results yielded a DEM set for each cohort (Figure 1E). In fusion-positive pRCC, 27 miRNAs were upregulated and 49 were downregulated, whereas 54 upregulated and 96 downregulated miRNAs were identified in the fusion-negative group. Further comparison identified 17 upregulated and 16 downregulated miRNAs specific to the fusion-positive cohort, and 44 upregulated and 63 downregulated miRNAs specific to the fusion-negative cohort (Figure 1E; Supplementary Table S4).

### Protein-Protein Interaction (PPI) network analysis

To further evaluate the functional relevance of the identified DEGs, protein–protein interaction (PPI) networks were constructed in both PRCC-TFE3 fusion-positive and fusion-negative KIRP cohorts. For the fusion-positive group, PPI networks were generated using the 880 upregulated and 301 downregulated exclusive DEGs. Three significant clusters of highly connected genes were identified among the upregulated DEGs, from which the ten most connected hub genes were RAB7A, CD8A, CDH1, GAPDH, CTSD, STAT1, SQSTM1, CCNB1, RACK1, and LAMP1. Among these, CD8A, STAT1, and LAMP1 were shared between the hub and module gene sets (Figure 2A–C). In the downregulated PPI network, the hub genes were TJP3, RAC3, DSP, KRT19, FGFR3, OCLN, CCND1, ACAA1, CLDN7, and SPINT1, with TJP3, RAC3, DSP, and OCLN shared between the hub and module groups (Figure 2B–C).

**Figure 2.**
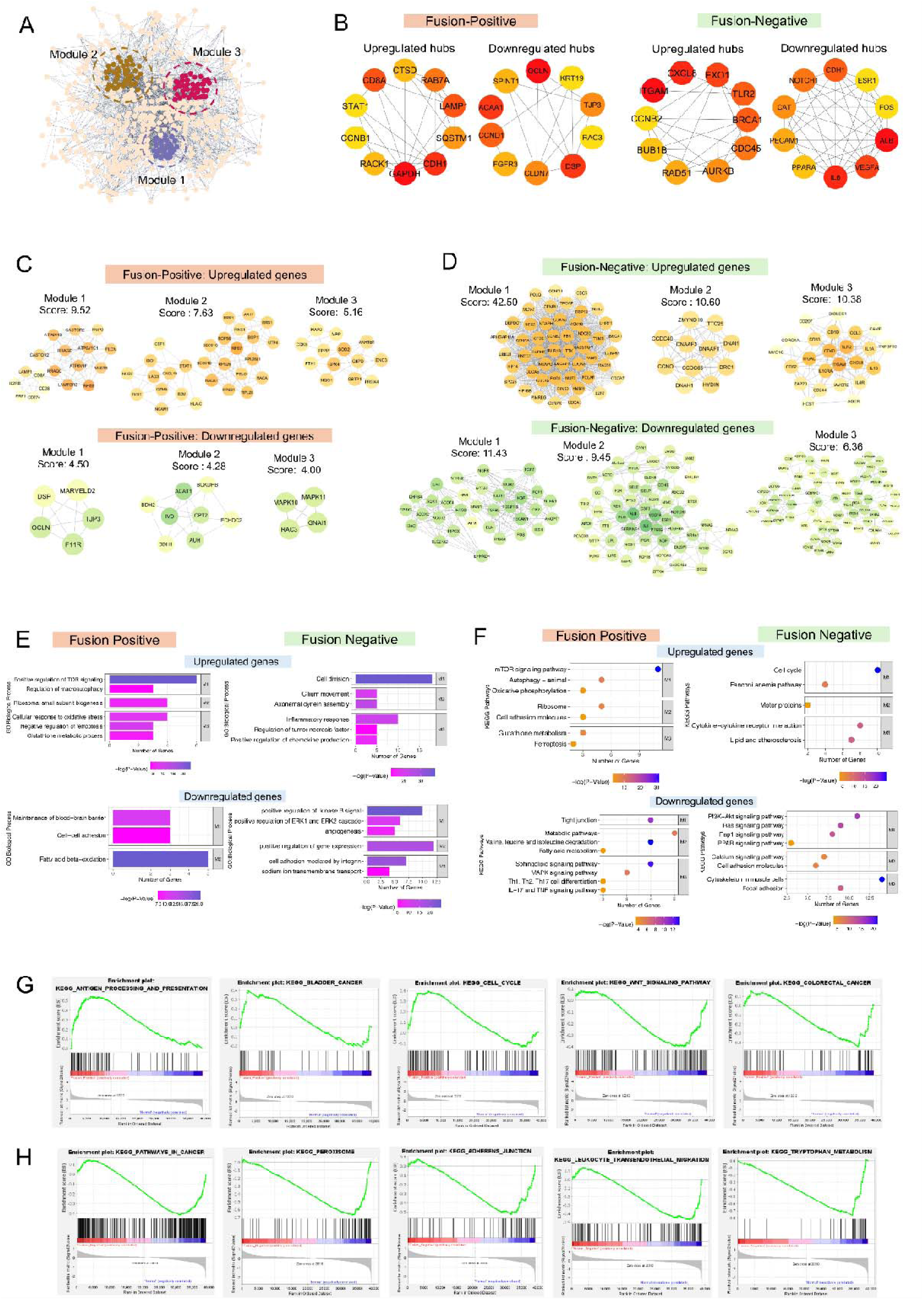
Comparative Protein-Protein Interaction Network and Enrichment Analyses in PRCC-TFE3 Fusion and Fusion-Negative KIRP. **A.** Schematic overview: PPI network-based modular clustering of DEGs in PRCC-TFE3 fusion-positive and fusion-negative KIRP cohorts, showing distinct modules (colored clusters) derived from exclusive upregulated or downregulated genes. **B.** Top ten upregulated and downregulated hub genes in each group ranked by CytoHubba scores, with a red-to-yellow color gradient indicating relative centrality. **C.** Fusion-positive cohort: modules derived from upregulated (top) and downregulated (bottom) DEGs, with node intensity reflecting degree of connectivity. **D.** Fusion-negative cohort: corresponding module structure and connectivity for up- and downregulated DEGs. **E–F.** Bar and bubble plots illustrating GO and KEGG enrichment for upregulated (top) and downregulated (bottom) DEGs from the major network modules. Circle size denotes gene counts; color gradient reflects p-value significance. **G–H.** GSEA plots showing significantly enriched KEGG pathways associated with (G) upregulated and (H) downregulated DEGs in fusion-positive (top row) and fusion-negative (bottom row) KIRP. Pathways with normalized enrichment scores (|NES| ≥ 1) are displayed; gene-rank positions indicate expression directionality.

In the fusion-negative cohort, PPI networks were constructed using 981 upregulated and 1,431 downregulated DEGs. The upregulated network yielded ten hub genes: CDC45, BRCA1, BUB1B, CXCL8, EXO1, TLR2, CCNB2, ITGAM, RAD51, and AURKB, while the downregulated network identified IL6, CAT, ALB, ESR1, CDH1, FOS, PPARA, PECAM1, VEGFA, and NOTCH1 as major hubs. All upregulated hub genes were contained within significant clusters, whereas nine downregulated hubs overlapped, with PPARA unique to the non-modular set (Figure 2B and D).

Collectively, these PPI networks indicate distinct interaction architectures in fusion-positive versus fusion-negative KIRP, providing insight into subtype-specific oncogenic circuits and highlighting candidate genes for downstream pathway and functional enrichment analyses.

### Differential pathway and functional enrichment in fusion-positive vs. fusion-negative KIRP

To evaluate the functional implications of differentially expressed genes (DEGs) specific to PRCC-TFE3 fusion status, we conducted enrichment analyses of significant network modules with emphasis on pathways associated with hub genes identified in fusion-positive and fusion-negative KIRP cohorts relative to normal kidney tissue (Figure 2E–F and Supplementary Table S1A).

### Fusion-Positive: transcriptional rewiring toward autophagy, metabolism, and mTOR signaling

In the fusion-positive group, significantly enriched biological processes were related to mTOR signaling activation, regulation of macroautophagy, ribosomal subunit biogenesis, negative regulation of ferroptosis, and oxidative stress response. These genes were associated with molecular functions including GTPase binding, cadherin and adaptor protein binding, and glutathione peroxidase activity, with subcellular localization predominantly at the plasma and lysosomal membranes (Figure 2E and Table S1B). KEGG pathway analysis of upregulated modules revealed enrichment in mTORC1 signaling (FLCN, RRAGC, FNIP2, MLST8), autophagy and phagosome pathways (LAMP1, ATP6V1C1, RRAGD), oxidative phosphorylation, ferroptosis and glutathione metabolism, supporting a reprogrammed metabolic phenotype (Figure 2F and Table S1A). LAMP1 was enriched in both the autophagy and phagosome pathways.

Conversely, downregulated DEGs in fusion-positive tumors were associated with blood–brain barrier-related functions, junctional integrity, and cell–cell adhesion, including reduced expression of tight junction proteins (e.g., OCLN, TJP3 and RAC3) (Figure 2E–F). These genes were also enriched in signaling pathways including MAPK, TNF, and IL-17/Th17 differentiation, along with alterations in fatty acid and butanoate metabolism, consistent with disrupted epithelial polarity and inflammatory signaling programs (Table S1B).

### Fusion-Negative: enrichment in cell cycle, inflammation, and adhesion pathways

In contrast, fusion-negative upregulated genes were primarily enriched in processes such as cell division, chemokine production, lipopolysaccharide response, and cell adhesion. Functional analysis indicated elevated serine/threonine kinase activity, ATPase activity, and cytokine signaling. Pathway enrichment indicated activation of the cell cycle, Fanconi anemia, NF-κB, TLR, and natural killer cell-mediated cytotoxicity pathways (Figure 2E–F; Table S1A–B). Upregulated hub genes, including CCNB2, BUB1B, CDC45, RAD51, CXCL8, and TLR2 were involved in these pathways. These findings support a hyperproliferative, pro-inflammatory state in the absence of TFE3 fusion.

Downregulated genes in fusion-negative tumors were associated with cytoskeletal remodeling, extracellular matrix (ECM) interactions, Ras/Rap1 signaling, PPAR signaling pathways, and cytoskeletal and cardiomyopathy-related pathways, consistent with loss of tissue structural homeostasis and altered metabolic programs (Figure 2E–F; Table S1A–B). CDH1, IL6, VEGFA, and NOTCH1 were among the downregulated hub genes linked to these pathways (Table S1A). Differential expression of DEGs within the identified modules for both fusion-positive and fusion-negative KIRP datasets is summarized in Table S2.

### Gene set enrichment analysis (GSEA) reveals divergent pathways by fusion status

GSEA further characterized distinct pathway activations linked to fusion status. In fusion-positive tumors, enriched pathways included antigen presentation and bladder cancer-related gene sets, with CTSD, LAMP1, CCNB1, CDH1 and CD8A as leading-edge contributors (Figure 2G). Notably, downregulation of Wnt signaling was a prominent feature of the fusion-positive cohort, with several key regulators, including CCND1, MAPK10, CTNNBIP1, and RAC3, selectively downregulated and overlapping with colorectal cancer pathway genes (Figure 2G).

In the fusion-negative cohort, GSEA indicated enrichment in peroxisome biogenesis, adherens junction formation, and leukocyte transendothelial migration, consistent with a more canonical pRCC signature (Figure 2H). Downregulated genes including NOTCH1, CDH1, and CAT contributed to these pathways, consistent with impaired cell–cell communication and oxidative defense.

### Network analysis of differentially expressed miRNAs and their gene targets

To examine the biological implications of fusion-specific miRNA profiles, putative targets of DEMs were identified. Predicted gene targets were compared with the differentially expressed genes (DEGs) identified in the corresponding cohorts to derive expression-concordant regulatory pairs, linking upregulated miRNAs to downregulated genes, and conversely.

In fusion-positive pRCC, the miRNA–mRNA interaction network included 967 pairs between 28 upregulated miRNAs and 236 downregulated genes, and 2,001 interactions between 28 downregulated miRNAs and 634 upregulated genes. Within this network, 14 downregulated gene targets were predicted to be regulated by 23 upregulated miRNAs, while 14 upregulated genes were predicted to be targeted by 15 downregulated miRNAs (Supplementary Figure S2).

In the fusion-negative pRCC cohort, 79 upregulated miRNAs were predicted to regulate 1,017 downregulated DEGs, and 110 downregulated miRNAs were predicted to regulate 1,411 upregulated DEGs. Module-associated genes were highlighted within the resulting network (Supplementary Figure S3), showing 108 downregulated module genes were predicted to be regulated by 76 upregulated miRNAs and 71 upregulated module genes were predicted to be regulated by 81 downregulated miRNAs.

### Downregulated miRNA activates translational, immune, redox, and mTOR pathways in PRCC-TFE3

To investigate post-transcriptional mechanisms associated with transcriptional activation in PRCC-TFE3 tumors, we mapped downregulated miRNAs to upregulated genes within the top PPI modules (Figure 3B). This analysis identified coordinated regulatory networks involving immune activation, oxidative metabolism, and lysosomal mTOR signaling. A redox-associated cluster comprising SOD2, GPX4, NQO1, and G6PD was predicted to be co-targeted by hsa-miR-874-5p, hsa-miR-30c-1/2-3p, hsa-miR-130a, and hsa-miR-497-5p, and SOD2 was predicted to be regulated by five distinct miRNAs. Binding energy profiles supported high-affinity interactions, consistent with functional derepression in oxidative stress response. hsa-miR-874-5p was identified as a broad-spectrum regulator, targeting genes across multiple modules including STAT1, SOD2, GPX4, ATP6V1C1, and NQO1, linking its downregulation to simultaneous immune activation, redox stability, and lysosomal acidification. Concurrently, miRNAs such as hsa-miR-27b-3p, hsa-miR-497-5p, hsa-miR-30c-1-3p, and hsa-miR-195-5p were predicted to regulate nutrient-responsive genes (FLCN, FNIP2, ATP6V1C1, LAMTOR2, MLST8) within the mTORC1 signaling axis, suggesting that loss of miRNA-mediated repression may facilitate metabolic reprogramming in fusion-positive PRCC. Collectively, these results indicate that loss of miRNA-mediated repression, particularly involving hsa-miR-874-5p, hsa-miR-30c-1/2-3p, and hsa-miR-497-5p, may permit the transcriptional activation of convergent oncogenic programs including mTOR signaling, immune modulation, and redox balance.

**Figure 3.**
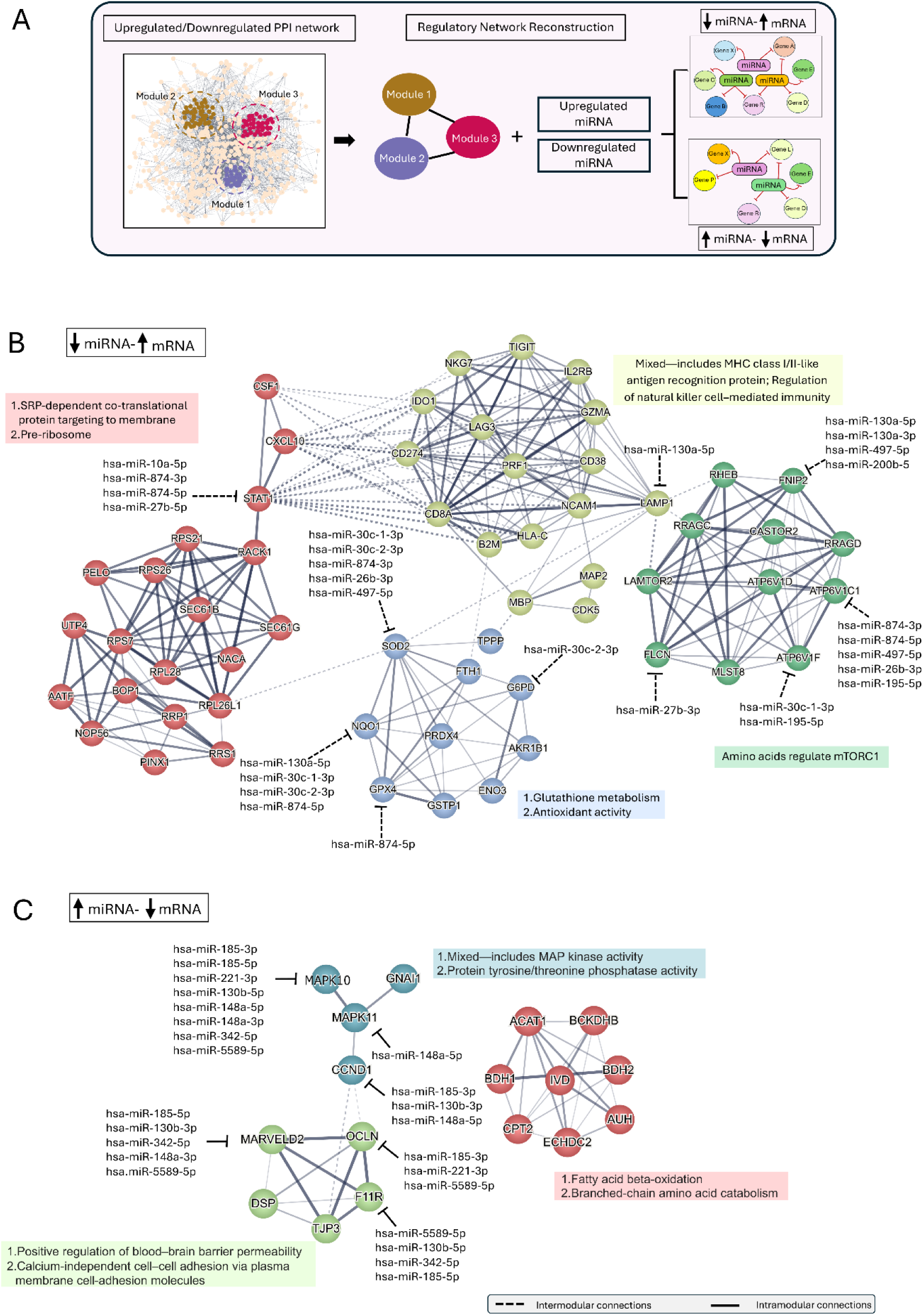
Schematic and functional mapping of miRNA-mRNA regulatory networks in PRCC-TFE3 fusion tumors. **A.** Schematic overview of the analytical workflow illustrating the reconstruction of regulatory networks using differentially expressed genes (DEGs) and miRNAs in PRCC-TFE3 tumors. The PPI network of upregulated and downregulated DEGs was partitioned into functional modules, followed by integration with upregulated and downregulated miRNAs to identify inverse miRNA–mRNA regulatory pairs. **B.** Network visualization of predicted interactions between downregulated miRNAs and upregulated mRNAs in the fusion-positive cohort. The network highlights modular regulation of translational machinery, redox metabolism, immune signaling, and mTORC1-associated nutrient sensing. Dashed lines denote intermodular connections, and solid lines indicate intramodular interactions. **C.** Network visualization of upregulated miRNAs targeting downregulated mRNAs in the fusion-positive cohort. Distinct modules include MAPK signaling, tight junction/cell adhesion, and fatty acid and branched-chain amino acid metabolism. Arrows represent miRNA-mediated repression. Shared miRNAs across modules suggest coordinated suppression of signaling and metabolic pathways.

### Upregulated miRNAs orchestrate suppression of key signaling, adhesion, and metabolic programs in Fusion-Positive PRCC-TFE3

To examine the role of upregulated miRNAs in modulating downregulated transcriptional programs in PRCC-TFE3 tumors, we reconstructed miRNA–mRNA regulatory networks from PPI modules enriched for downregulated genes (Figure 3C). A MAPK signaling cluster comprising MAPK10, MAPK11, GNAI1, and CCND1 was predicted to be suppressed by hsa-miR-185-3p, hsa-miR-130b-5p, hsa-miR-148a-5p, hsa-miR-221-3p, hsa-miR-342-5p, and hsa-miR-5589-5p, with MAPK10 as a central node. These interactions indicate that elevated miRNA expression may dampen MAP kinase pathway activity in fusion-positive tumors. Tight junction and adhesion components including MARVELD2, OCLN, F11R, TJP3, and DSP formed a second module, predicted to be targeted by hsa-miR-185-5p, hsa-miR-130b-3p, hsa-miR-342-5p, hsa-miR-148a-3p, hsa-miR-221-3p, and hsa-miR-5589-5p. MARVELD2 and F11R were commonly targeted, consistent with disruption of epithelial cohesion. Notably, hsa-miR-185-5p, hsa-miR-130b-3p, and hsa-miR-148a-3p were shared across modules, suggesting that they act as central regulators suppressing signaling, structural, and metabolic programs in fusion-positive PRCC. These results highlight the integrative role of miRNA networks in shaping the transcriptional landscape of PRCC-TFE3 tumors through modular, multitarget regulation.

### Structural validation of miRNA-mRNA duplexes highlights AGO2-compatible targeting in PRCC-TFE3

Structural validation of miRNA–mRNA duplexes supports AGO2-compatible targeting in PRCC-TFE3. To validate the regulatory interactions associated with transcriptomic alterations in PRCC-TFE3 tumors, we performed RNAhybrid-based prediction of miRNA–mRNA duplexes and assessed their structural compatibility with AGO2 loading using AlphaFold 3 (Figure 4 and Supplementary Figure S5A). Among the predicted interactions, several miRNA–mRNA pairs displayed minimum free energy values below −25 kcal/mol and formed ternary AGO2 complexes with high model confidence (ipTM ≥ 0.82, pTM ≥ 0.90) (Figure 4A and B).

**Figure 4.**
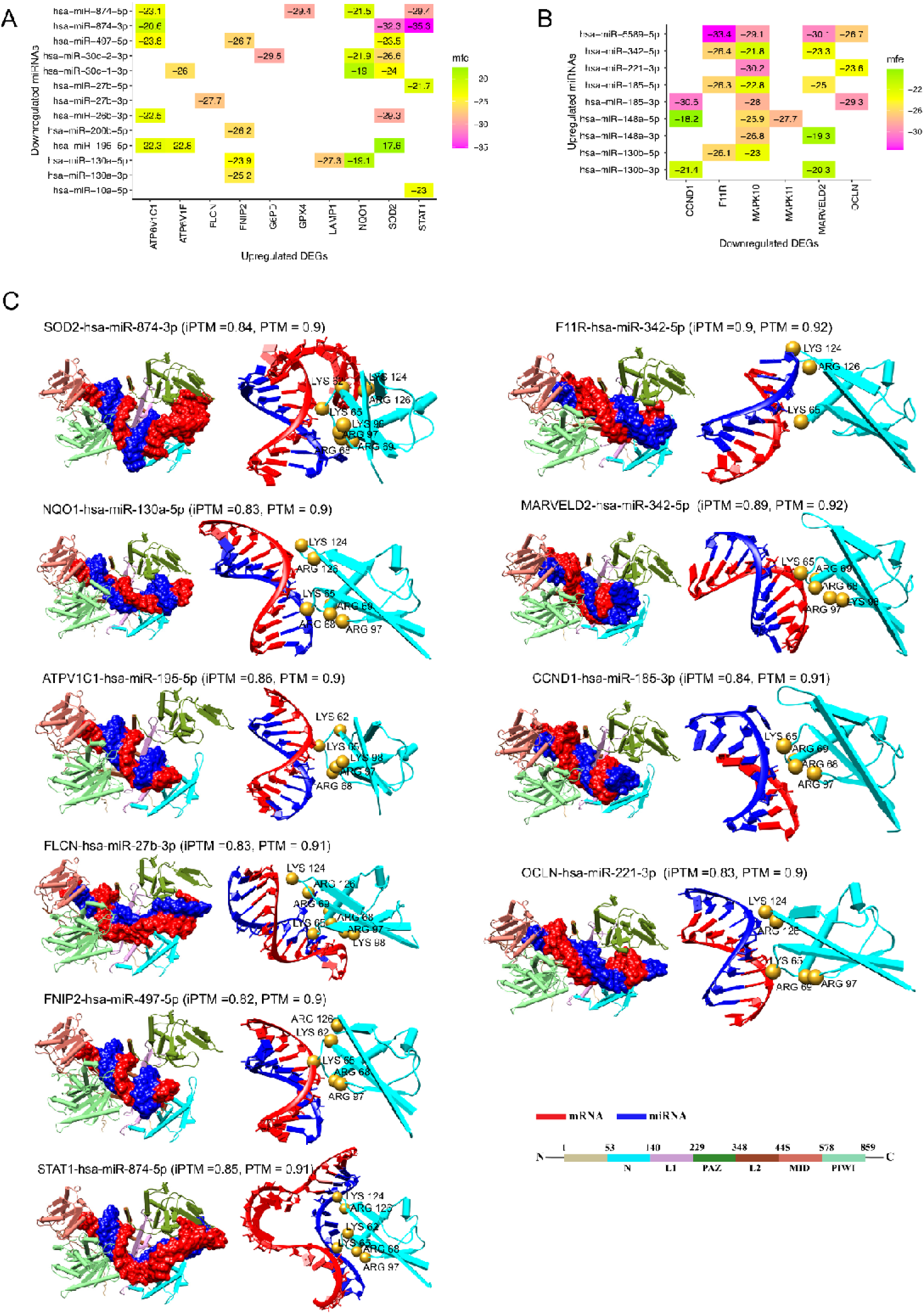
Structural modeling of miRNA–mRNA duplexes and AGO2 interactions in PRCC-TFE3 tumors. A–B. RNAhybrid-predicted minimum free energy (MFE) values of top-ranked miRNA–mRNA interactions involving upregulated genes predicted to be targeted by downregulated miRNAs **(A)** and downregulated genes predicted to be targeted by upregulated miRNAs **(B)** in fusion-positive PRCC. **C.** Ternary complex models of selected duplexes docked to human AGO2, generated using AlphaFold 3. Each panel shows HsAGO2 domains, mRNA (red), and miRNA (blue). Key AGO2 residues (Lys62, Lys65, Arg68, Arg69, Arg97, Lys98, Lys124, Arg126) that interact with the duplex backbone are highlighted. ipTM and pTM scores denote model confidence for protein–RNA complexes.

Within the fusion-positive cohort, several duplexes linked to antioxidant, mTOR, and immune signaling pathways showed stable placement within the central AGO2 channel. These included SOD2–hsa-miR-874-3p (ipTM 0.84), NQO1–hsa-miR-130a-5p (ipTM 0.83), ATP6V1C1–hsa-miR-195-5p (ipTM 0.86), FLCN–hsa-miR-27b-3p (ipTM 0.83), FNIP2–hsa-miR-497-5p (ipTM 0.82), and STAT1–hsa-miR-874-5p (ipTM 0.85). These miRNA–mRNA duplexes occupied the central groove of AGO2, stabilized through electrostatic interactions with positively charged residues at the N-domain interface, including Lys62, Lys65, Arg68, Arg69, Arg97, Lys98, Lys124, and Arg126, consistent with RISC-compatible configurations [37] (Figure 4C).

In the fusion-positive downregulated network, stable duplexes were observed for tight junction and metabolic targets. These included F11R–hsa-miR-342-5p (ipTM 0.90), MARVELD2–hsa-miR-342-5p (ipTM 0.89), CCND1–hsa-miR-185-3p (ipTM 0.84), and OCLN–hsa-miR-221-3p (ipTM 0.83). These complexes showed canonical docking within AGO2, supporting their potential relevance in suppressing adhesion and signaling pathways in fusion-negative tumors (Figure 4C). The structural evidence supports the regulatory plausibility of selected miRNA–mRNA interactions in mediating fusion-context–specific post-transcriptional repression in PRCC.

For the fusion-negative cohort, RNAhybrid identified 25 high-affinity miRNA–mRNA pairs (Supplementary Figure S5B). AlphaFold 3 modeling produced seven duplexes with ipTM ≥ 0.8 and pTM ≥ 0.9 (Supplementary Table S6). Six of these—BRCA1–hsa-miR-660-3p, CCNB2–hsa-miR-483-5p, ESR1–hsa-miR-6892-5p, BRCA1–hsa-miR-135a-2-3p, FOS–hsa-miR-1295a, and CCNB2–hsa-miR-654-5p—showed AGO2-compatible conformations similar to those seen in fusion-positive models (Figure 5).

**Figure 5.**
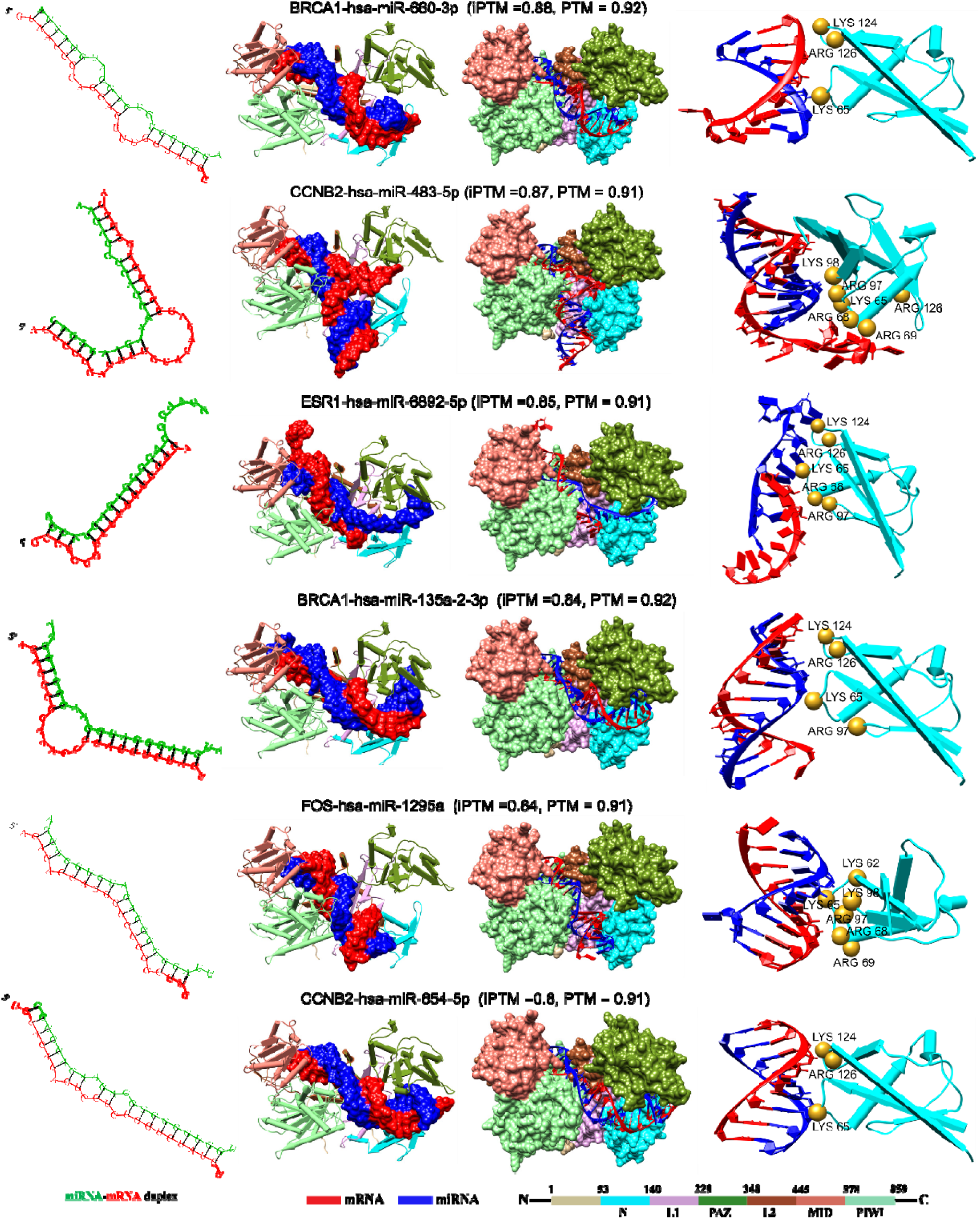
Structural modeling of miRNA–mRNA duplexes and AGO2 interactions in fusion-negative KIRP cases. RNAhybrid-predicted high-affinity miRNA–mRNA duplex pairs are shown in the left panel. AI-based structural models of selected duplexes docked to human AGO2 were generated with AlphaFold 3. Each panel shows HsAGO2 domains (cyan), mRNA (red), and miRNA (blue). Key AGO2 residues interacting with the duplex backbone are highlighted. ipTM and pTM scores denote model confidence for protein–RNA complexes (right panel).Identification of PRCC- and TFE3-targeting miRNAs and their differentially expressed gene targets

To assess downstream consequences of PRCC 3′UTR deletion and TFE3 fusion transcript formation, we analyzed a subset of 15 miRNAs predicted to bind PRCC and shown to be expressed in kidney tissue (Table S5 and Figure S4A). Network reconstruction identified 745 miRNA–gene regulatory pairs between these PRCC-targeting miRNAs and 236 downregulated genes, primarily mapping onto enriched modules identified in fusion-positive pRCC (Figure S4A). Notably, only hsa-miR-3074-5p was also upregulated among differentially expressed miRNAs, suggesting it may contribute to gene repression in fusion tumors. The network showed that these PRCC-targeting miRNAs were also predicted to regulate hub downregulated genes across modules involved in signaling (e.g., MAPK1, RAC3), metabolic processing (e.g., BDH1, ACAD8), adhesion (e.g., CLDN7, OCLN, F11R), and key regulators, including GNAI1, CCND1, and MARVELD2. These genes are also predicted targets of upregulated DEMs and were supported by three-dimensional modeling, supporting their potential functional relevance. Importantly, the pattern of interaction suggests a redistribution of miRNA targeting: in the absence of wild-type PRCC 3′UTR, several miRNAs may redirect binding activity toward alternative targets, amplifying repression of signaling, structural, and metabolic genes.

To further evaluate the post-transcriptional effects of fusion-driven miRNAs, we mapped upregulated miRNAs predicted to target the TFE3 transcript to downregulated DEGs in fusion-positive PRCC. Among 122 predicted TFE3-interacting miRNAs, five were significantly upregulated in tumor samples: hsa-miR-197-5p, hsa-miR-185-3p, hsa-miR-185-5p, hsa-miR-342-5p, and hsa-miR-221-3p. These miRNAs formed a regulatory network with 183 downregulated DEGs, yielding 540 miRNA–mRNA interactions (Figure S4B and Supplementary Table S5). Notably, several genes central to tight junctions, cell adhesion, and signaling, including OCLN, F11R, MARVELD2, MAPK10, MAPK11, GNAI1, and DSP, were predicted to be co-targeted by both TFE3- and PRCC-binding miRNAs. Although PRCC-targeting miRNAs were not differentially expressed in tumors, TFE3-associated miRNAs (e.g., hsa-miR-185-3p, hsa-miR-342-5p, hsa-miR-148a-5p, hsa-miR-130b-5p, hsa-miR-221-3p) showed increased expression, suggesting they may potentially substitute for miRNAs lost with PRCC 3′UTR truncation (Table S3). For instance, hsa-miR-342-5p and hsa-miR-185-3p simultaneously targeted MARVELD2, OCLN, F11R, and DSP, consistent with coordinated repression of epithelial adhesion integrity. MAPK10 and MAPK11, regulators of MAPK signaling cascades, were also targeted by both hsa-miR-185-3p and hsa-miR-197-5p, implicating miRNA-mediated dampening of stress-responsive pathways.

### Immune cell infiltration patterns in Fusion-Positive and Fusion-Negative PRCC-TFE3

Because several of the identified genes mapped to immune-related pathways, immune infiltration patterns were evaluated between fusion-positive and fusion-negative cases from the TCGA-KIRP cohort. The ESTIMATE algorithm was used to estimate the immune score, stromal score, and combined ESTIMATE score for each sample. Immune and ESTIMATE scores were significantly higher in fusion-positive pRCC than in fusion-negative tumors and normal tissues (Figure 6). The MCP-counter algorithm was used to estimate the relative abundance of nine immune and stromal cell populations. Fusion-positive tumors exhibited lower inferred abundance of CD3⁺ T cells and neutrophils, whereas CD8⁺ T cells, macrophages, endothelial cells, and natural killer (NK) cells were comparatively higher. The proportions of B cells, myeloid dendritic cells, and cancer-associated fibroblasts (CAFs) were similar between the two groups, consistent with distinct immune infiltration profiles across fusion states.

**Figure 6.**
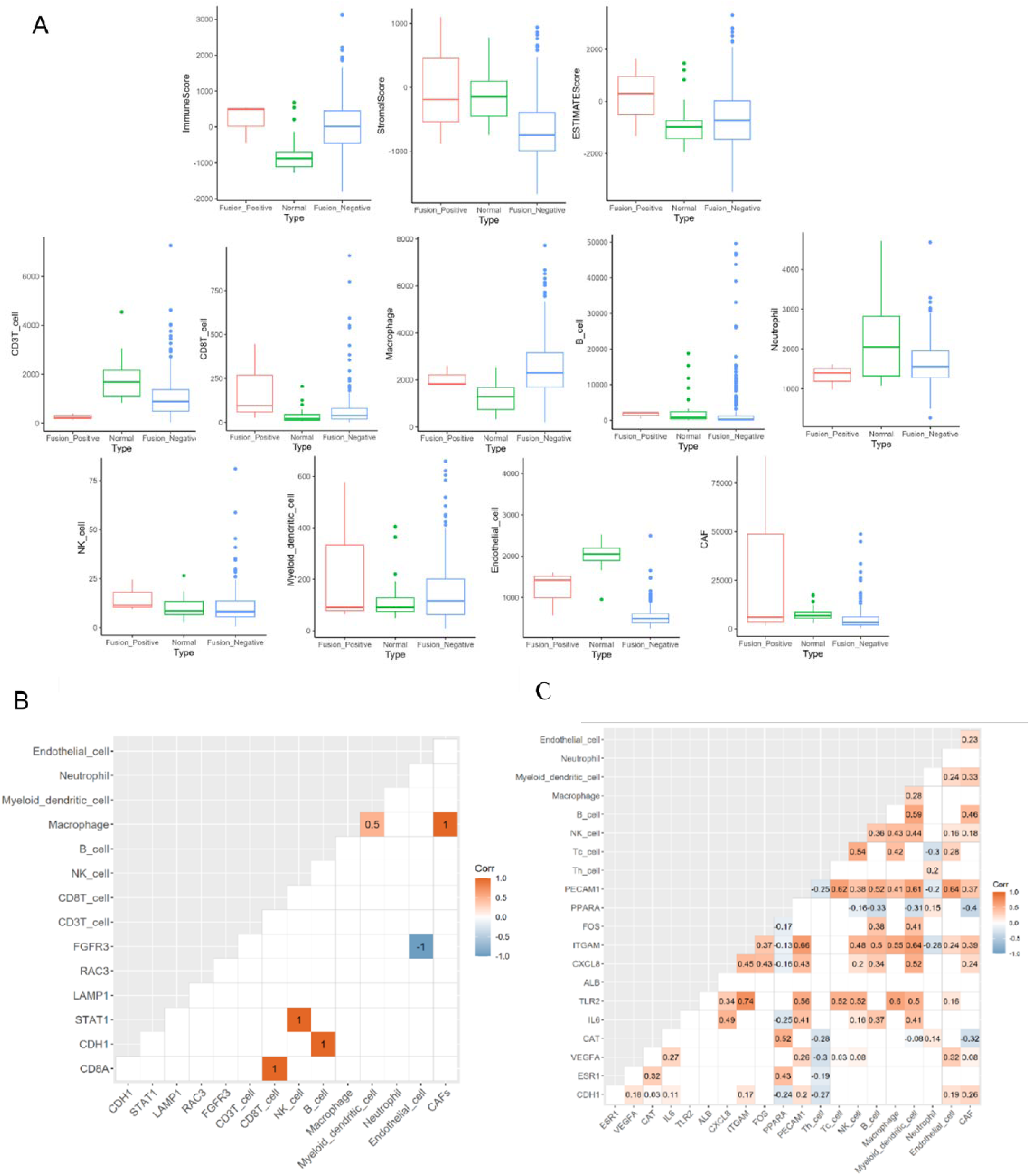
Comparison of immune microenvironments between fusion-positive and fusion-negative KIRP cases. **(A)**The ESTIMATE algorithm reveals significantly higher immune, stromal, and ESTIMATE scores in fusion-positive KIRP than in fusion-negative KIRP. MCP-counter revealed differential infiltration levels of immune and stromal cell types between fusion-positive and fusion-negative KIRP cohorts. Correlation plot of immune-related hub genes and immune cells in **(B)** fusion-positive KIRP and **(C)** fusion-negative KIRP.

To further examine immune-related genes, immune-associated hub genes were identified in both groups (Figure 6B–C). In fusion-positive pRCC, positive correlations were observed between CD8⁺ T cells and CD8A, NK cells and STAT1, and B cells and CDH1, whereas the downregulated gene FGFR3 showed a negative correlation with endothelial cell abundance. Positive associations were also observed among macrophages, myeloid dendritic cells, and CAFs. In fusion-negative PRCC, correlation analysis between immune-related genes and immune cell types revealed broader and more complex associations compared with the fusion-positive context. Positive correlations were evident between TLR2 and multiple immune cell types, including macrophages (0.60), NK cells (0.52), myeloid dendritic cells (0.50), and Tc cells (0.52), consistent with a role in immune interactions. CXCL8 showed a positive association with myeloid-derived cells (dendritic cells) (0.50), supporting its pro-inflammatory function. PECAM1 correlated positively with endothelial cells (0.64), Tc cells (0.62), and myeloid dendritic cells (0.61). Overall, fusion-negative PRCC exhibits more distributed immune gene–cell correlations, consistent with a more heterogeneous immune microenvironment compared with fusion-positive tumors (Figure 6C).

### ROC analysis of candidate biomarkers

Receiver Operating Characteristic (ROC) analysis was conducted for 17 genes (11 upregulated and 6 downregulated) identified in fusion-positive pRCC. All genes showed AUC ≥ 0.7. RRAGC showed a maximum AUC = 1.0, followed by ATP6V1F, F11R, FNIP2, GPX4, MAPK11, MARVELD2, OCLN, and RRAGD with AUC ≈ 0.99, indicating strong discrimination for fusion-positive samples (Figure 7A). For miRNAs, 10 candidates were evaluated; hsa-miR-130a showed the highest discriminative performance (AUC = 0.99), followed by hsa-miR-148a, hsa-miR-221, hsa-miR-342, hsa-miR-30c-2, hsa-miR-874, hsa-miR-130b, and hsa-miR-30c-1 (AUC > 0.90). hsa-miR-26b and hsa-miR-5589 showed AUC = 0.84 and 0.79, respectively (Figure 7B). In the fusion-negative group, PECAM1 showed the highest predictive performance (AUC = 0.97). Additional genes with AUC ≥ 0.90 included AURKB, CDC45, CAT, ALB, CDH1, VEGFA, and NOTCH1 (Figure 7C). For miRNAs, hsa-miR-654 and hsa-miR-650 showed the highest AUC values (0.91), followed by hsa-miR-483 (0.89), hsa-miR-370 (0.87), hsa-miR-135a-2 (0.85), and hsa-miR-1293 (0.80) (Figure 7D).

**Figure 7.**
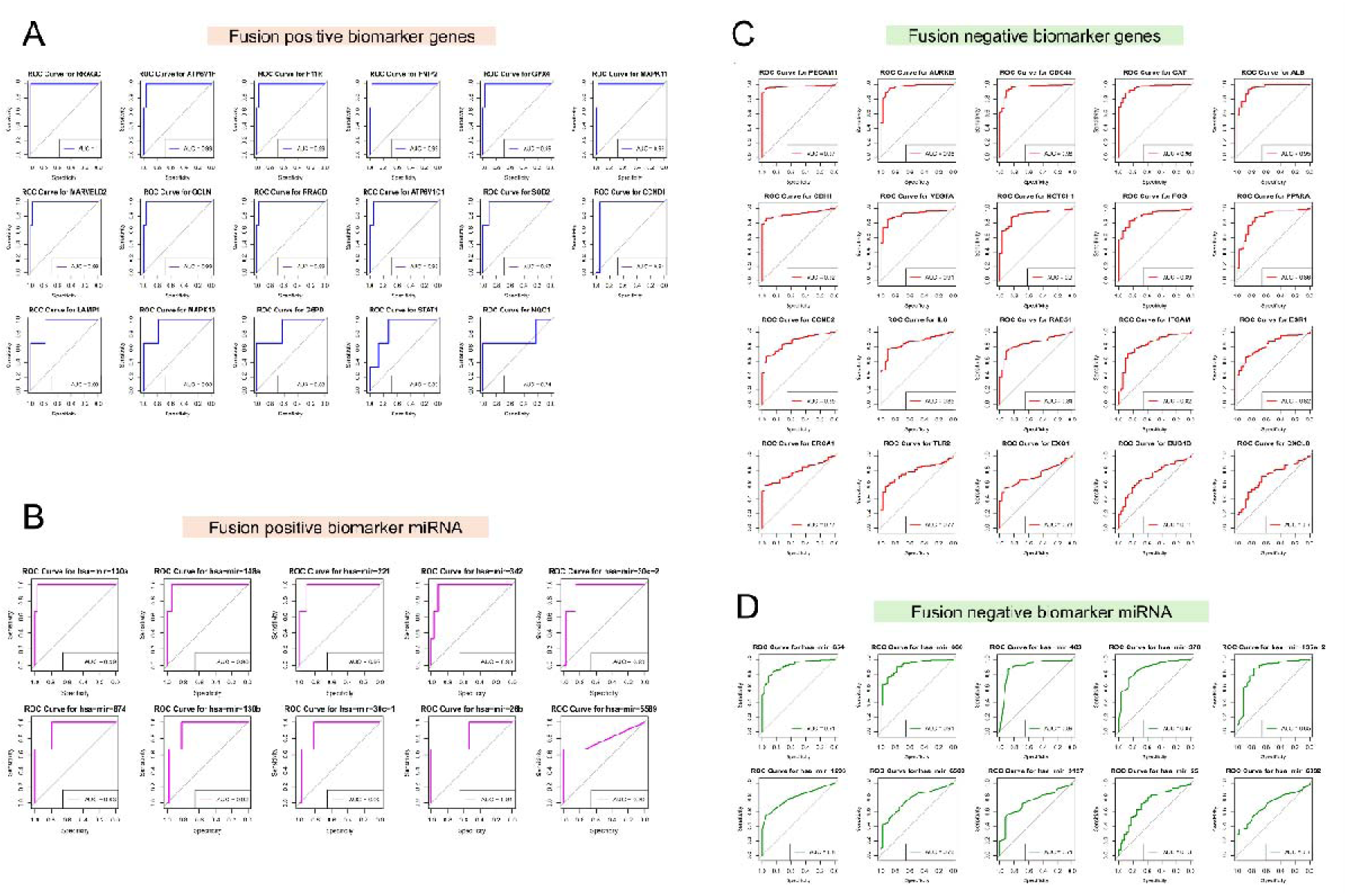
ROC curves of identified biomarker genes and miRNAs in fusion-positive and fusion-negative KIRP cases. The figure shows AUC values and ROC curves for **(A)** fusion-positive KIRP genes and **(B)** fusion-positive KIRP miRNAs, and **(C)** fusion-negative KIRP genes and (D) fusion-negative KIRP miRNAs. AUC ≥ 0.7 was used as the performance threshold.

### Connectivity Map (CMap) Analysis

Connectivity Map analysis was used to identify small molecules with gene-expression signatures inversely correlated with the DEG signatures. Compounds were ranked by normalized connectivity scores (Table 1). One compound was prioritized for the fusion-positive cohort and three for the fusion-negative group. In fusion-positive pRCC, TAK-715, a p38-MAPK inhibitor, was predicted to reverse the expression signature, consistent with dysregulation of MAPK10/11 in this study. For fusion-negative pRCC, three compounds—cladribine, fenretinide, and mozavaptan—were prioritized, targeting ADA, CYP3A5, and AVPR1B, respectively, each of which was upregulated in fusion-negative samples.

**Table 1.** List of drugs obtained from CMap analysis based on normalised connectivity score, mechanism of action and drug targets.

| Pert_iname | MOA | Targeted_upregulated_degs | Norm_cs |
| --- | --- | --- | --- |
| **Fusion Positive** |  |  |  |
| TAK-715 | P38 MAPK inhibitor | – | -1.73 |
| **Fusion Negative** |  |  |  |
| Cladribine | Adenosine deaminase inhibitor,<br>Ribonucleoside reductase inhibitor | ADA | -1.95 |
| Fenretinide | Apoptosis stimulant, Retinoid receptor<br>agonist | CYP3A5 | -1.92 |
| Mozavaptan | Vasopressin receptor antagonist | AVPR1B | -1.9 |

## Discussion

TFE3-rearranged papillary renal cell carcinoma is a biologically aggressive variant of pRCC, yet the mechanisms through which PRCC-TFE3 fusions reshape tumor behavior have largely been interpreted through transcriptional alterations and metabolic pathway activation [8, 75–81]. By integrating matched mRNA–miRNA sequencing, network inference, and AGO2-based structural modeling, this study defines a fusion-specific post-transcriptional architecture that links ferroptosis resistance, metabolic rewiring, epithelial adhesion loss, dormancy signatures, and immune remodeling in PRCC-TFE3 tumors. These results refine current models of fusion-driven oncogenesis by indicating that fusion-induced transcriptional programs are closely coupled to miRNA-mediated derepression and AGO2-compatible regulatory complexes, thereby suggesting a multi-layered RNA-dependent regulatory system.

Consistent with recent proteogenomic and transcriptomic analyses of TFE3-rearranged RCC [8, 81], fusion-positive tumors were enriched for oxidative phosphorylation, mTOR signaling, and glutathione metabolism (Figure 2). Within this signature, the coordinated upregulation of SOD2, GPX4, G6PD, and NQO1 was particularly notable, as these genes constitute a redox homeostasis and ferroptosis-suppressive program. These enzymes are key regulators of oxidative stress, supporting mitochondrial homeostasis. SOD2 detoxifies mitochondrial superoxide; GPX4 eliminates phospholipid hydroperoxides; G6PD provides NADPH essential for glutathione recycling; and NQO1 modulates ROS levels while coupling redox flux to autophagy–mTOR pathways [82–88]. Collectively, these activities define a ferroptosis-resistant metabolic state that is consistent with the high OXPHOS dependency reported in TFE3-driven renal carcinomas.

A key advance of this study is delineation of the miRNA circuitry that enables this metabolic rewiring. Integration of miRNA–mRNA pairing identified a redox-centric cluster including SOD2, GPX4, G6PD, and NQO1 that is typically constrained by miRNAs such as hsa-miR-874-5p, hsa-miR-30c-1-3p, hsa-miR-30c-2-3p and hsa-miR-497-5p (Figure 3B). Downregulation of these miRNAs in fusion-positive tumors thus removes a critical post-transcriptional checkpoint, enabling increased transcriptional output of antioxidant programs. Among these, hsa-miR-874-5p emerged as a broad-spectrum miRNA, with predicted targets including STAT1, SOD2, GPX4, ATP6V1C1, and NQO1, suggesting that its suppression may concurrently reinforce immune signaling, oxidative stress tolerance, and lysosomal acidification. Similar regulatory patterns were observed for hsa-miR-30c family and hsa-miR-221, hsa-miR-342, and hsa-miR-130b, which have been implicated in renal cancer biology and prognostic stratification [89–94]. Concomitant deregulation of nutrient-sensing miRNAs including hsa-miR-27b-3p, hsa-miR-497-5p, and hsa-miR-195-5p further relieved repression of the FLCN–FNIP2–LAMTOR–mTORC1 signaling axis. These miRNAs typically constrain mTOR activation by targeting FLCN, FNIP2, ATP6V1C1, LAMTOR2, and MLST8, and their decreased expression in fusion-positive tumors may facilitate sustained mTORC1 engagement and anabolic remodeling (Figure 3B).

Beyond redox regulation, the miRNA–mRNA regulatory axis identified in fusion-positive tumors also extends to signaling, adhesion, and metabolic programs that collectively align with a dormancy-competent tumor state. Upregulated miRNAs, including hsa-miR-185-3p/5p, hsa-miR-130b-5p/3p, hsa-miR-148a-5p/3p, hsa-miR-221-3p, hsa-miR-342-5p, and hsa-miR-5589-5p, were predicted to downregulate key components of Wnt and MAPK signaling, notably MAPK10, MAPK11, GNAI1, and CCND1 (Figure 3C). This coordinated suppression of proliferative and stress-responsive cascades is consistent with prior observations that Wnt–MAPK attenuation facilitates cellular quiescence and metastatic dormancy [95, 96]. In keeping with this, tight junction genes central to epithelial barrier integrity, including MARVELD2, OCLN, F11R, TJP3, and DSP, were predicted to be targeted by the same set of upregulated miRNAs, suggesting disruption of epithelial adhesion architecture through post-transcriptional repression.

Downregulation of mitochondrial metabolic genes such as BDH1, ACAT1, BCKDHB, IVD, AUH, and CPT2, many of which were predicted targets of PRCC-associated miRNAs, further supports a low-metabolic and low-proliferative phenotype characteristic of dormant tumor states.

Several of the upregulated miRNAs identified here have reported functional relevance in renal and solid tumor biology: hsa-miR-185 has been linked to EMT and proliferative control in renal disease [97]; hsa-miR-148a has been reported to modulate Wnt-dependent growth programs in renal carcinoma [98]; hsa-miR-130b has been reported to be dysregulated in kidney-associated inflammatory conditions [99]; hsa-miR-221 has been associated with poor survival in RCC [100, 101]; and hsa-miR-342-5p has been proposed as a circulating biomarker in other epithelial malignancies [102, 103]. Their coordinated elevation in fusion-positive tumors thus aligns with established oncogenic or modulatory roles, yet here they appear reoriented toward promoting adhesion loss, MAPK suppression, and metabolic suppression.

Notably, multiple adhesion and signaling genes including OCLN, F11R, MARVELD2, MAPK10, MAPK11, GNAI1, and DSP were predicted to be co-targeted by both TFE3- and PRCC-interacting miRNAs. This convergence indicates that the fusion event not only introduces new miRNA-binding contexts on the chimeric transcript but also redistributes miRNA targeting toward shared downstream nodes (Figure S4). Two miRNAs upregulated in fusion-positive tumors, hsa-miR-342-5p and hsa-miR-185-3p, simultaneously targeted MARVELD2, OCLN, F11R, and DSP, consistent with disruption of epithelial cohesion. Likewise, the simultaneous targeting of MAPK10 and MAPK11 by hsa-miR-185-3p and hsa-miR-197-5p suggests miRNA-mediated attenuation of stress-activated MAP kinase signaling. Together, these patterns indicate a fusion-driven recalibration of post-transcriptional control in which TFE3-interacting miRNAs augment and may partially substitute for the regulatory functions lost due to PRCC 3′UTR truncation.

This integrated transcriptional and post-transcriptional remodeling supports a dual adaptive program in fusion-positive pRCC: activation of antioxidant and lysosomal–mTOR circuits that support ferroptosis resistance, coupled with miRNA-dependent repression of metabolic, proliferative, and junctional pathways that support cellular dormancy and immune evasion. This bifurcated regulatory architecture provides a mechanistic basis for the persistent, metastatic-competent, yet slow-cycling behavior characteristic of PRCC-TFE3 tumors.

Thermodynamic and structural analyses provide additional support for these interactions at the level of individual complexes. RNAhybrid predicted high-affinity duplexes (MFE < −18 kcal/mol), while AlphaFold 3 modeling demonstrated that several miRNA–mRNA pairs including SOD2–hsa-miR-874-3p, NQO1–hsa-miR-130a-5p, ATP6V1C1–hsa-miR-195-5p, FLCN–hsa-miR-27b-3p, FNIP2–hsa-miR-497-5p, and STAT1–hsa-miR-874-5p adopt AGO2-compatible conformations (Figure 4). These duplexes occupied canonical AGO2 binding grooves, stabilized by electrostatic complementarity, supporting post-transcriptional gene silencing. Notably, these structural observations parallel recent reports that TFE3 fusion proteins engage RNA-mediated condensates to amplify transcriptional output [104]. In this work, AGO2-bound miRNA complexes represent a complementary post-transcriptional layer that influences transcript stability and turnover, supporting a dual RNA-dependent regulatory logic in TFE3-driven tumor evolution.

The tumor immune microenvironment (TIME) influences metabolic adaptation, dormancy, and therapeutic response in solid tumors [105–107]. Immune infiltration analysis of the TCGA-KIRP cohort indicated that PRCC-TFE3 fusion-positive tumors exhibit a distinct immune landscape characterized by reduced CD3⁺ T-cell and neutrophil infiltration signals, yet increased inferred abundance of CD8⁺ T cells, NK cells, macrophages, and endothelial cells (Figure 6). Diminished CD3⁺ T-cell abundance, given their central role in TCR-dependent antitumor cytotoxicity, is consistent with impaired immune surveillance and is consistent with the aggressive clinical behavior described for TFE3-rearranged renal cancers [105, 108–110].

The reduced neutrophils further reflects an altered metabolic niche. Fusion-positive tumors showed upregulation of oxidative phosphorylation and a concomitant repression of fatty-acid metabolic genes (Figure 2), a configuration that mirrors contexts in which neutrophils adopt fatty-acid oxidation-driven immunosuppressive programming [111–113]. Although CD8⁺ T cells and NK cells were enriched, such increases have previously been associated with exhausted or dysfunctional effector states in renal malignancies, especially in metabolically constrained microenvironments [114, 115]. These combined shifts point not to global immune suppression, but to immune remodeling that favors tumor persistence. This immune profile is consistent with the broader dormancy-competent phenotype emerging from our transcriptomic and miRNA analyses. Reduced MAPK signaling, attenuated proliferative pathways, and enhanced autophagy-all detected in fusion-positive tumors-mirror hallmarks of survival-oriented dormancy programs in which immune modulation and reduced metabolic demand coexist to sustain long-term tumor viability and immune evasion (Figure 2, Figure 3).

The reduced neutrophils may reflect an altered metabolic niche. Fusion-positive tumors showed upregulation of oxidative phosphorylation with concomitant repression of fatty-acid metabolic genes (Figure 2), a configuration consistent with contexts in which neutrophils adopt fatty-acid oxidation-driven immunosuppressive programming [111–113]. Although CD8⁺ T cells and NK cells were increased, such increases have been associated with exhausted or dysfunctional effector states in renal malignancies, especially in metabolically constrained microenvironments [114, 115]. These combined shifts suggest not global immune suppression, but immune remodeling that may favor tumor persistence. This immune profile is consistent with the broader dormancy-competent phenotype identified from our transcriptomic and miRNA analyses. Reduced MAPK signaling, attenuated proliferative pathways, and enhanced autophagy—all detected in fusion-positive tumors—are consistent with hallmarks of survival-oriented dormancy states in which immune modulation and reduced metabolic demand coexist to sustain long-term tumor viability and immune escape (Figure 2, Figure 3).

To assess biomarker potential, ROC analyses identified a panel of genes that distinguish fusion-positive from fusion-negative tumors. In fusion-positive pRCC, genes associated with mTOR signaling (RRAGC, RRAGD, FNIP2, ATP6V1F, ATP6V1C1), ferroptosis resistance (GPX4, SOD2, G6PD, NQO1), immune signaling (STAT1), and epithelial integrity (F11R, OCLN) each showed AUC ≥ 0.7 (Figure 7). Together, these pathways define the fusion-positive transcriptional program mapped here. Conversely, fusion-negative tumors were distinguished by markers of proliferative cycling (AURKB, CDC45, BUB1B, RAD51), inflammatory signaling (CXCL8, TLR2, IL6), and vascular/metabolic regulation (VEGFA, PPARA, ALB, ESR1), consistent with a distinct biological profile.

To prioritize therapeutic candidates predicted to reverse these transcriptomic signatures, Connectivity Map (CMap) analysis was used on fusion-stratified DEG sets. In fusion-positive tumors, TAK-715, a p38-MAPK inhibitor with reported cross-reactivity against casein kinase I isoforms involved in Wnt regulation, showed strong negative connectivity, consistent with our observation that MAPK10/11 repression and Wnt pathway attenuation are defining features of this subtype [116]. For fusion-negative tumors, three compounds (cladribine, fenretinide, and mozavaptan) were prioritized as candidates. Their reported mechanisms including purine analog-induced lymphocyte apoptosis (cladribine) [117], retinoid-mediated autophagic cell death (fenretinide) [118], and vasopressin-receptor blockade (mozavaptan) [119] were consistent with distinct metabolic and receptor-level dysregulation identified in the fusion-negative profile.

Although this study uses one of the few available matched miRNA–mRNA datasets for PRCC-TFE3, several limitations remain. The rarity of fusion-positive cases in TCGA limits cohort size. Immune infiltration were inferred computationally and would benefit from validation by spatial or single-cell profiling in future studies. Finally, although structural modeling suggests AGO2-competent miRNA–mRNA duplex formation, biochemical confirmation in TFE3 fusion backgrounds remains necessary. Despite these limitations, the use of matched transcriptomic and miRNA sequencing, coupled with network and structural interrogation, provides a rigorous systems-level framework for dissecting the regulatory complexity of fusion-driven renal cancers.

In summary, this study delineates a two-layer regulatory program in PRCC-TFE3 fusion-positive pRCC in which transcriptional activation of oxidative and lysosomal pathways is reinforced through miRNA-mediated derepression of ferroptosis-resistant genes, while upregulated miRNAs suppress adhesion, MAPK, Wnt, and metabolic pathways to support a dormancy-compatible phenotype. AGO2-bound structural modeling supports the regulatory role of key miRNA–mRNA interactions, and immune deconvolution indicates a remodeled TIME that is consistent with the dormancy and metabolic signatures. The identification of fusion-specific biomarkers and the prioritization of TAK-715 as a therapeutic candidate support the translational relevance of these findings and provide a framework for precision approaches targeting TFE3-rearranged renal cancers.

## Materials and Methods

### Data retrieval, dimensionality reduction and data preprocessing

RNA-Seq and miRNA-Seq data of papillary renal cell carcinoma (pRCC) were downloaded from the TCGA-KIRP cohort through the GDC portal (https://portal.gdc.cancer.gov) [38], and PRCC-TFE3 fusion-positive cases were annotated using ChimerDB 4.0 [39]. To assess transcriptional differences between tumor and adjacent normal tissue in papillary renal cell carcinoma (KIRP), we analyzed RNA-seq data from three PRCC-TFE3 fusion-positive tumors, 287 fusion-negative KIRP tumors, and 32 normal kidney samples.

Dimensionality reduction using Principal Component Analysis (PCA) and Multidimensional Scaling (MDS) was used to examine sample distribution, identify potential outliers, and classify fusion-positive, fusion-negative, and normal samples [40, 41]. PCA of the fusion-positive group showed clear separation between tumor and normal samples, with PC1 and PC2 explaining 43.5% and 14.5% of the total variance, respectively. The fusion-negative cohort also showed clear clustering, with PC1 and PC2 explaining 31% and 8.8% of the variance (Supplementary Figure S1). MDS supported these patterns, with separation evident along the first two components (35% and 24% variance in fusion-positive; 15% and 8% in fusion-negative), supporting a distinct transcriptomic separation between tumor and normal tissues.

### Differential Gene and miRNA expression analyses

Differential expression analyses were performed using the DESeq2 and LIMMA packages in R to identify differentially expressed genes (DEGs) and differentially expressed miRNAs (DEMs), using thresholds of adjusted p-value (padj) < 0.05 and |log2FC| ≥ 1 for the PRCC-TFE3 fusion-positive vs. normal and fusion-negative vs. normal comparisons [42, 43]. Volcano plots were produced using the EnhancedVolcano R package (version 4.5.0) [44], and Venn diagrams were generated using the InteractiVenn web tool [45].

### Protein-Protein Interaction (PPI) network

PPI networks for DEGs were constructed using the STRING application in Cytoscape v3.9.1. [46, 47]. Hub genes were identified using CytoHubba [48], and functional modules were identified using MCODE (degree cutoff = 2, node score cutoff = 0.2, K-core = 2, max depth = 100) [49].

### Functional and pathway enrichment analysis

Functional and pathway enrichment analyses of fusion-positive and fusion-negative TCGA-KIRP samples were conducted using the DAVID tool [50–52]. Gene Ontology (GO) terms and pathways with p < 0.05 were considered significant and visualized with ggplot2 in R [53]. Gene set enrichment analysis (GSEA) was performed using the c2.cp.kegg.v2023.1.Hs.symbols.gmt collection from MSigDB with the GSEA software (version 4.3.2), using default settings [54, 55].

### MicroRNA gene target prediction and DEM-DET regulatory network analysis

MicroRNA expression data were retrieved from TCGA, and mature miRNA sequences were obtained from the miRBase database (https://www.mirbase.org/). Target prediction for DEMs was conducted using the miRWalk database [56]. miRNAs predicted to target the PRCC gene, together with their putative targets, were extracted from miRWalk. Only targets with predicted 3′UTR binding sites were retained for further analysis. We compared miRNA gene targets with DEGs to identify differentially expressed targets (DETs) in PRCC-TFE3 fusion-positive and fusion-negative cases using the InteractiVenn web tool. Downregulated gene targets of PRCC- and TFE3-targeting miRNAs, exclusive to fusion-positive pRCC, were then identified.

Regulatory networks linking upregulated DEMs to their downregulated target genes, and downregulated DEMs to their upregulated targets, were constructed in Cytoscape v3.9.1. Based on these networks, the top 10 hub miRNAs targeting PRCC were identified based on degree centrality. The miRNA–mRNA regulatory network was refined to emphasize hub DEMs and hub DEGs that also appeared in the PPI network. In parallel, we examined the regulatory network associated with PRCC- and TFE3-targeting miRNAs and their corresponding DETs. From this analysis, we selected hub miRNAs targeting both PRCC and TFE3, also based on degree centrality. Finally, interaction pairs comprising these hub miRNAs and their corresponding downregulated hub genes from the PPI network were delineated.

### Prediction and validation of miRNA-mRNA interactions

Mature miRNA sequences were obtained from the miRBase database (https://www.mirbase.org), and 3′ UTR regions of target mRNAs were obtained from UTRdb v2.0 [57]. In silico interaction screening was conducted using RNAhybrid [58] to assess binding between differentially expressed miRNAs (DEMs) and their differentially expressed target genes. miRNA–mRNA pairs with minimum free energy (mfe) ≤ −18 kcal/mol were retained as candidate interactions. Interaction profiles were visualized as heatmaps using the pheatmap package in R [59].

### Structural modeling of miRNA-mRNA duplexes and Argonaute 2 complexes

Secondary structures of selected miRNA–mRNA duplexes were predicted using RNAhybrid [60]. To model AGO2-compatible conformations, the sequence of human Argonaute 2 (HsAGO2) was obtained from the UniProt database [61]. Argonaute proteins typically bind guide RNAs to form RNA-induced silencing complexes (RISCs) that mediate endonucleolytic cleavage of complementary transcripts [37, 62].

Three-dimensional models of miRNA–mRNA duplexes bound to HsAGO2 were generated using AlphaFold 3 [63]. Predicted complexes were ranked based on interface predicted template modeling (ipTM) and predicted template modeling (pTM) scores. Thresholds of ipTM ≥ 0.8 and pTM ≥ 0.9 were applied to support high confidence in relative interface positioning and overall structural accuracy. Default random seeds were used, and the highest-ranked structures were carried forward for downstream analysis. Structural interactions of the miRNA–mRNA–HsAGO2 complexes were analyzed and visualized using UCSF Chimera v1.18 [64].

### Estimation of tumor purity and microenvironmental cell infiltration

Tumor purity, stromal scores, and immune scores were estimated for fusion-positive and fusion-negative TCGA-KIRP samples with the ESTIMATE R package [65]. The infiltration of stromal and immune cell populations, including total T cells, CD8⁺ T cells, cytotoxic cells, NK cells, B cells, monocytes, macrophages/monocytes, myeloid dendritic cells, neutrophils, endothelial cells, and cancer-associated fibroblasts, was estimated using the Microenvironment Cell Populations-counter (MCP-counter) algorithm as implemented in the Immunedeconv R package [66, 67].

### Retrieval of immune related genes

Immune-related genes were obtained from the Immunome and ImmPort databases [68, 69]. Differentially expressed immune-related hub genes were identified through comparative analysis between fusion-positive and fusion-negative TCGA-KIRP cohorts.

### Correlation of immune cell infiltration with immune hub genes

Spearman’s correlation analysis was used to assess associations between immune cell infiltration levels and immune-related hub gene expression within the fusion-positive and fusion-negative TCGA-KIRP cohorts. Correlation matrices were generated using the corrplot R package [70], with statistical significance defined as correlation coefficient > 0.7 and p < 0.05.

### Receiver operating characteristic (ROC) analysis

Genes and miRNAs prioritized from PPI and functional analyses were evaluated using ROC curve analysis. Normalized expression values were derived using the estimateSizeFactors function in R [71]. ROC curves and area under the curve (AUC) values were computed with the pROC package [72], and curves were visualized using the plotROC function in ggplot2 [73]. Predictive performance was considered acceptable when AUC ≥ 0.7.

### Connectivity map (CMap) analysis

The CMap platform [74] was used to prioritize small molecules with potential therapeutic relevance. For both fusion-positive and fusion-negative cohorts, upregulated DEGs from modules 1–3 were combined into the upregulated query set, and downregulated DEGs into the downregulated query set. Queries were run against the Touchstone repository (L1000 gene expression signatures). Compounds were ranked by normalized connectivity scores, where positive scores indicate concordant effects whereas negative scores indicate inhibitory (reversal) effects. The top 20 negative-score compounds were retained, excluding compounds with unknown mechanisms of action. Drug targets were annotated, and only compounds targeting genes significantly upregulated in our dataset or pathways significantly dysregulated were considered.

## Declaration of competing interest

The authors declare that there are no conflicts of interest with the contents of this article.

## CRediT Authorship Contribution Statement

D M: methodology, software, data curation, formal analysis, visualization, writing - original draft, review & editing. S A: software, formal analysis, investigation, visualization, writing - review & editing. Divya: software, writing - review & editing. E P: conceptualization, formal analysis, visualization, investigation, methodology, writing - review & editing. RM: conceptualization, methodology, investigation, writing - original draft, writing - review & editing, visualization, supervision, project administration funding acquisition.

## Acknowledgments

We gratefully acknowledge the following sources of support: D.M. for JRF and SRF from DBT Govt. of India; S.A. for project assistant support from DST-CURIE-2022-80, Government of India; Divya for CSIR-JRF and SRF; R.M. gratefully acknowledges for the research support from DST-CURIE-2022-80(G), Government of India.

## Notes

### Competing Interest Statement

The authors have declared no competing interest.

