## Supplementary_Material for "Fusion-driven post-transcriptional network orchestrates ferroptosis resistance, dormancy, and immune remodeling in PRCC-TFE3 renal cell carcinoma"

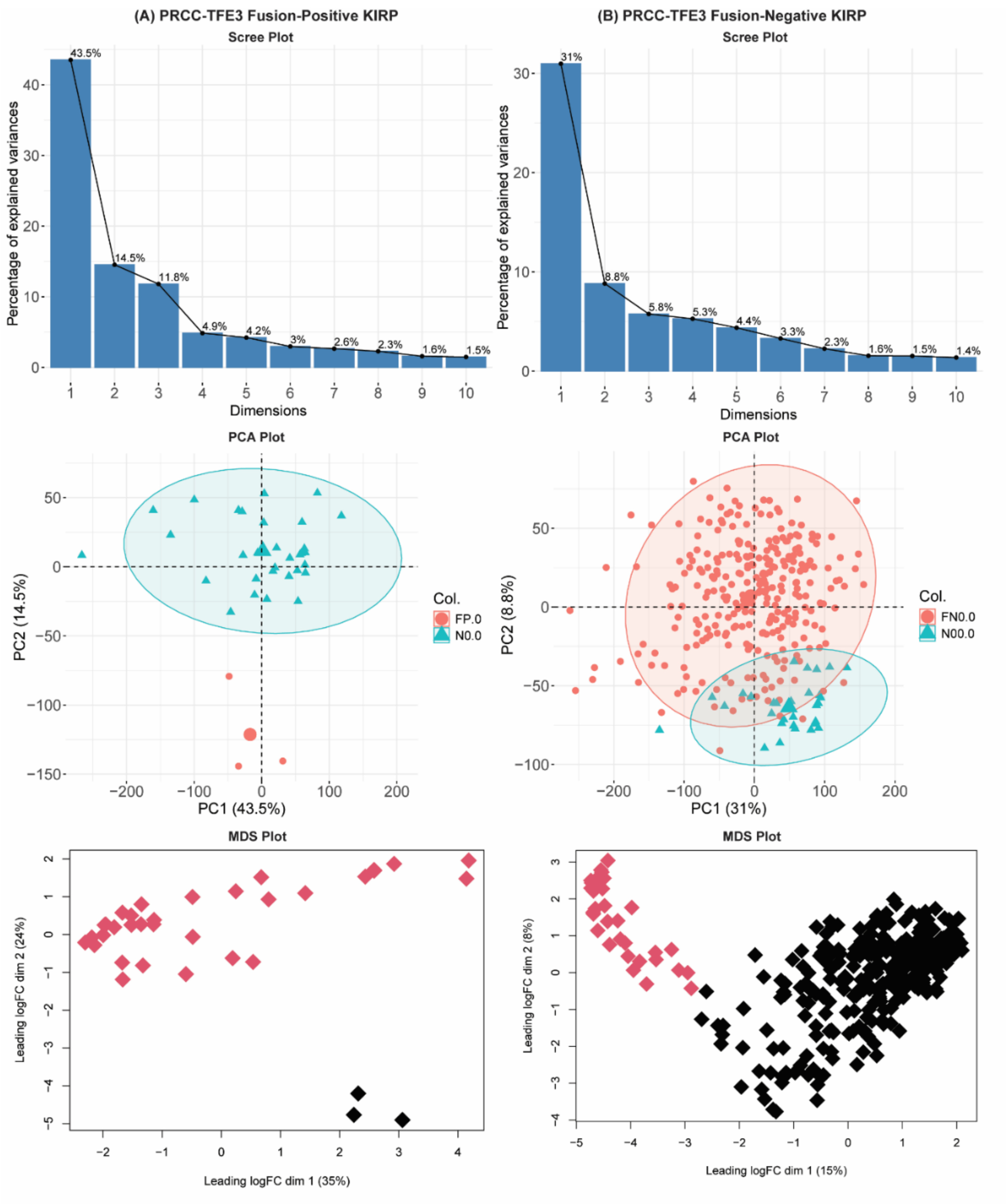


**Supplementary Figure S1.** Principal component analysis (PCA) and Multidimensional scaling (MDS) analysis of the (A) Fusion-positive vs normal and (B) Fusion-negative vs normal datasets. Scree plot showing possible numbers of PCs might be calculated in the dataset. We have selected PC1 and PC2 for each dataset for pair wise PCA plot generation and cluster analysis. MDS plot shows variation among samples based on normalized data. The distance between each sample shows the dissimilarity between those two samples. Red colored diamond shows normal and black colored diamond shows tumor samples.

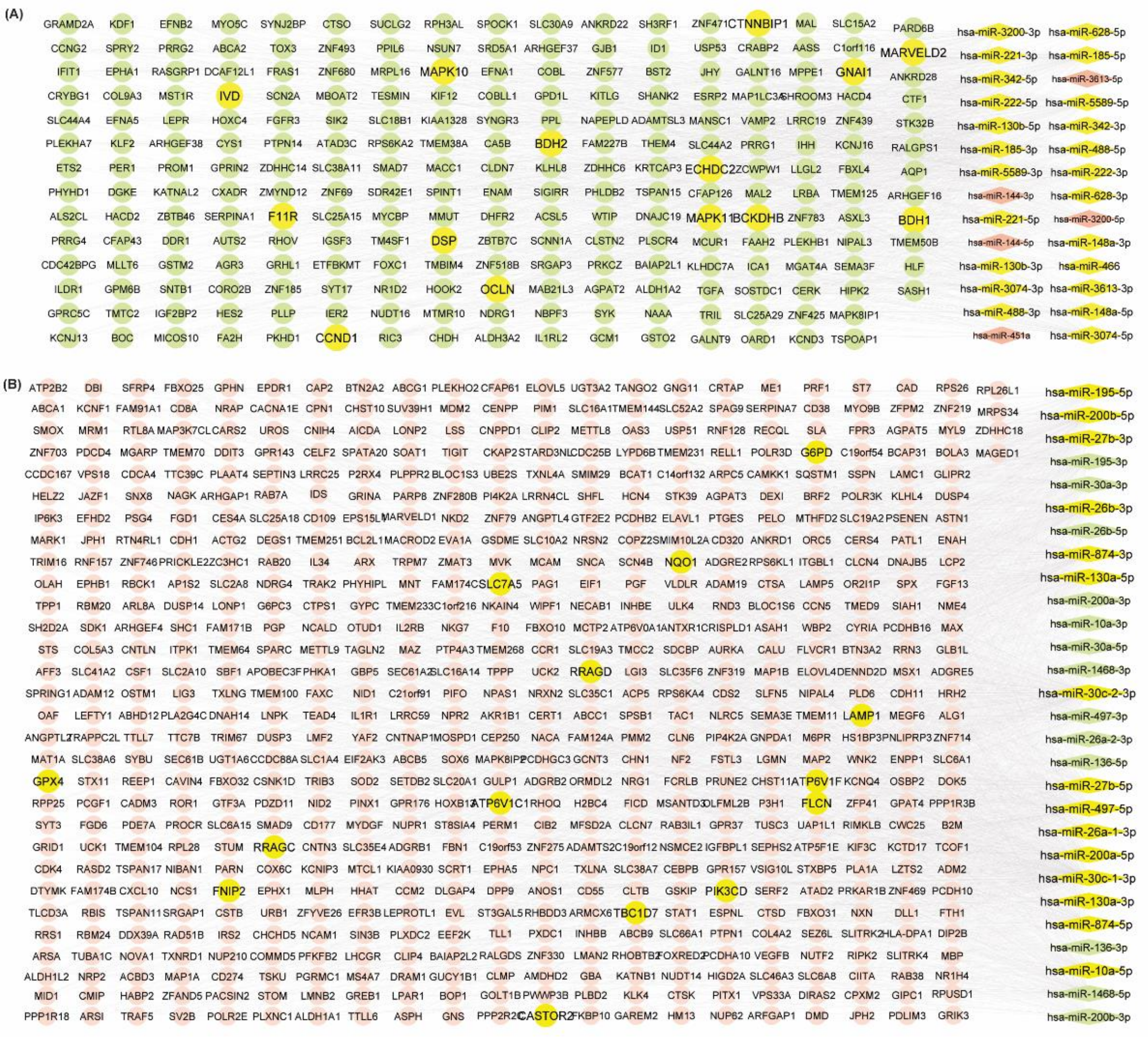

**Supplementary Figure S2.** The miRNA-gene regulatory network depicts the differential interaction between DEMs and their gene targets. The interactions are shown between the A. Downregulated targets of the upregulated DEMs in fusion-positive KIRP B. Upregulated targets of the downregulated DEMs in fusion-positive KIRP. Yellow nodes are the genes of modules targeted by the differentially expressed miRNAs. Red color represents upregulation and green color represents downregulation of the genes and miRNAs. Differentially expressed targets are shown using circular shaped nodes and DEMs are shown using diamond shaped nodes.

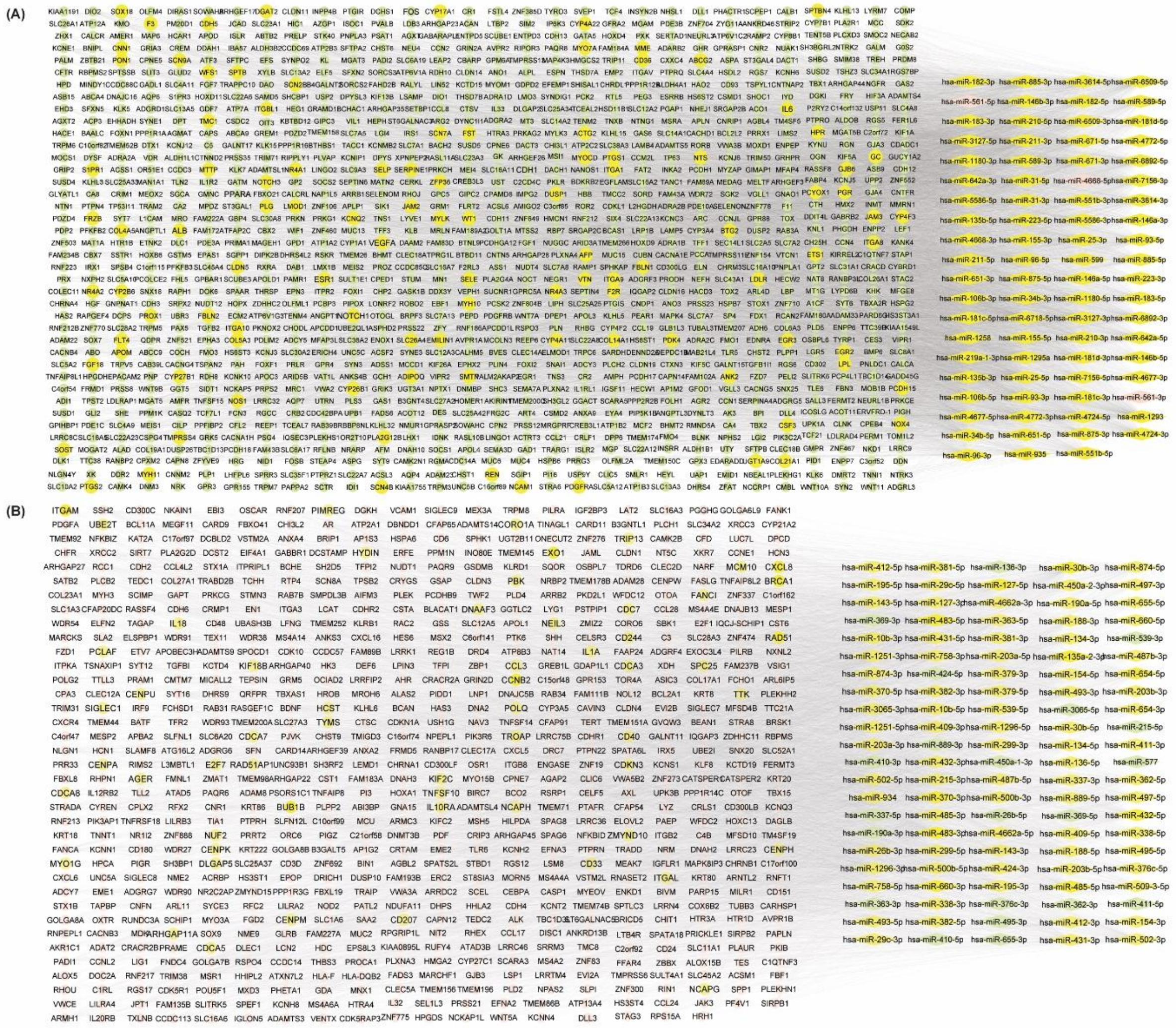

**Supplementary Figure S3.** The miRNA-gene regulatory network depicts the differential interaction between DEMs and their gene targets. The interactions are shown between the A. Downregulated targets of the upregulated DEMs in fusion-negative KIRP B. Upregulated targets of the downregulated DEMs in fusion-negative KIRP. Yellow nodes are the genes of modules targeted by the differentially expressed miRNAs. Red color represents upregulation and green color represents downregulation of the genes and miRNAs. Differentially expressed targets are shown using circular shaped nodes and DEMs are shown using diamond shaped nodes.

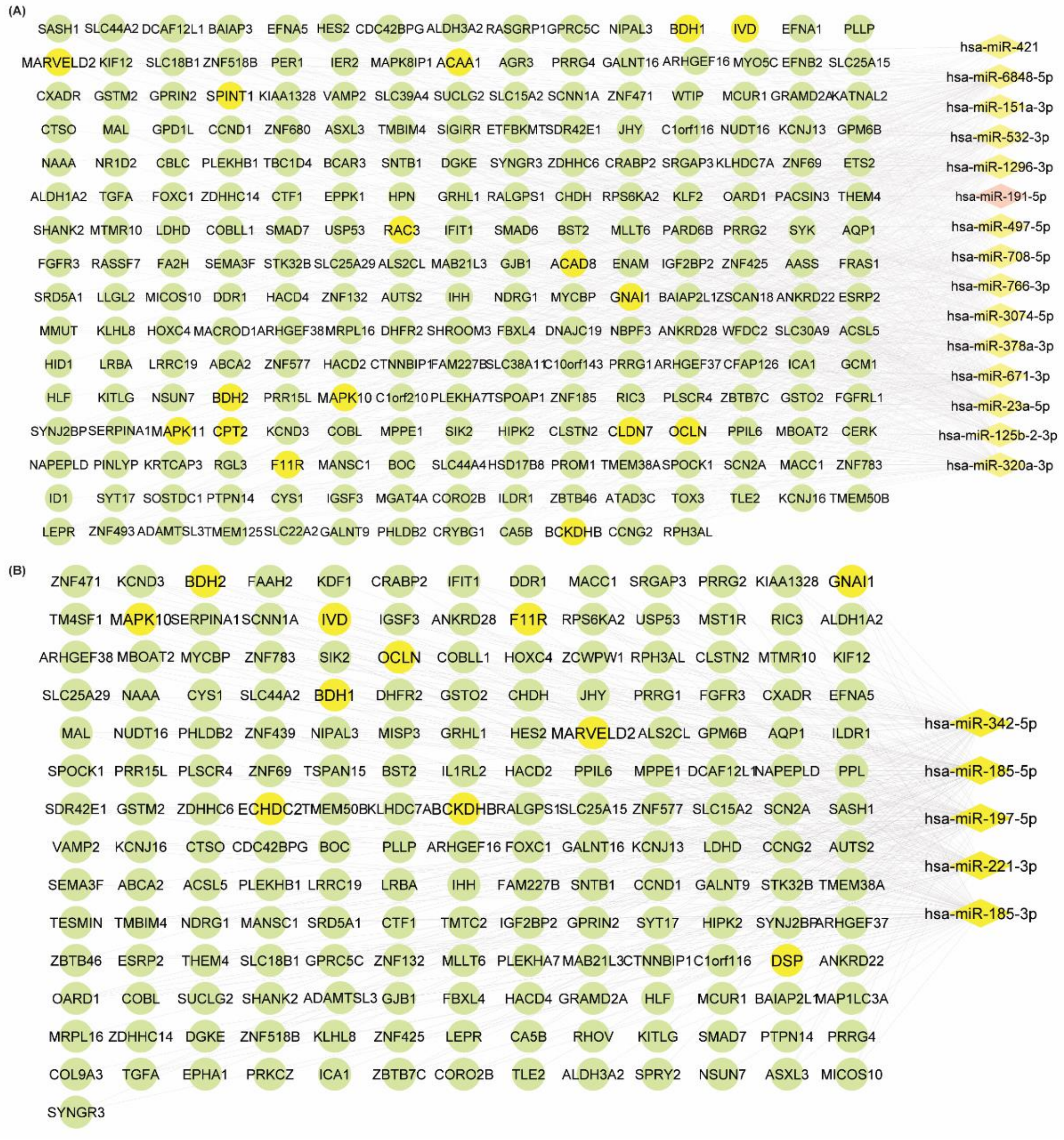


**Supplementary Figure S4.** miRNA–mRNA regulatory network illustrating interactions between A) PRCC-targeting miRNAs and downregulated DEGs in fusion-positive pRCC. B) TFE3-targeting upregulated miRNAs showing differential targeting with their downregulated target DEGs. Yellow circles represent downregulated genes from key modules, and yellow diamonds represent miRNAs previously predicted to target PRCC and expressed in kidney tissue. Edges represent predicted regulatory interactions. The observed network highlights broad and overlapping repression by PRCC-targeting miRNAs on genes involved in signaling, metabolism, and adhesion.

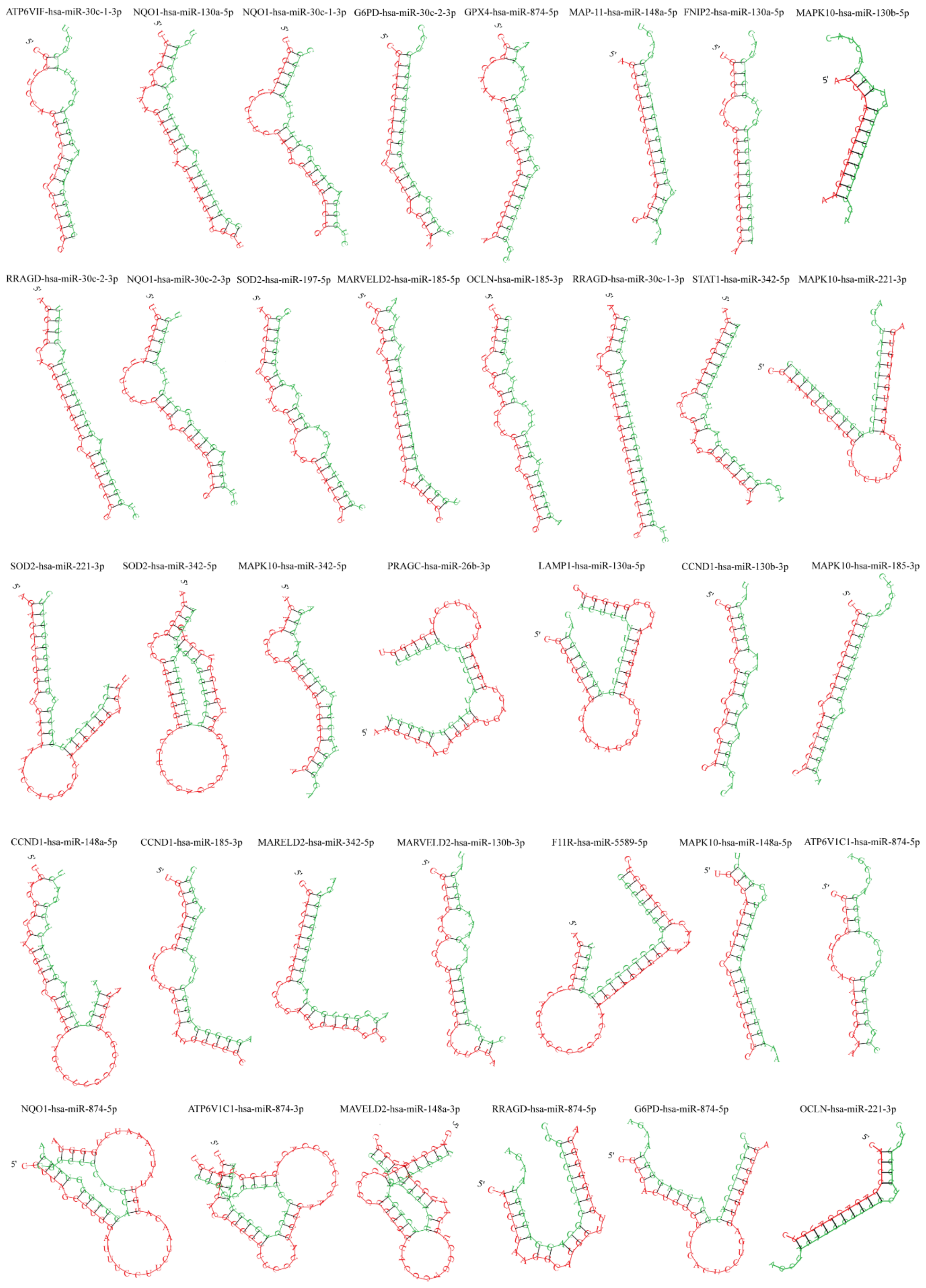


**Supplementary figure S5A.** 2D structures of influential miRNA-mRNA pairs in fusion positive KIRP cases.

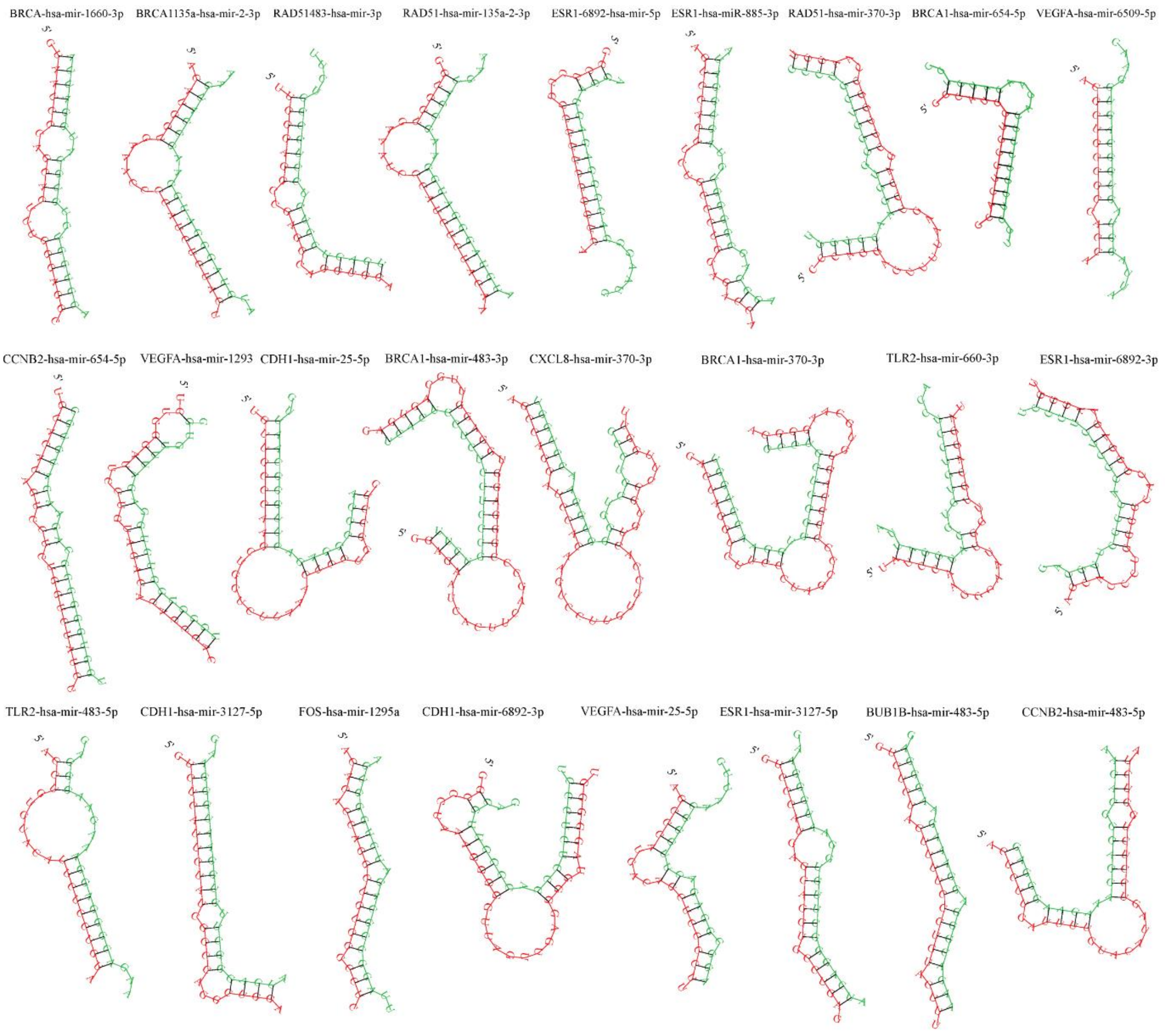


**Supplementary figure S5B.** 2D structures of influential miRNA-mRNA pairs in fusion negative KIRP cases.

**Supplementary Table S1A.** List of significant pathways linked to modules of fusion positive and negative cases.

| Modules | Term | Genes (*hub gene) | PValue |
|---|---|---|---|
| | **Fusion-Positive Vs Normal: Upregulated DEGs** | | |
| **Module 1** | hsa04150:mTOR signaling pathway | FLCN, RRAGC, CASTOR2, RHEB, RRAGD, MLST8, LAMTOR2, FNIP2, ATP6V1D, ATP6V1C1, ATP6V1F | 1.83E-14 |
| | hsa04140:Autophagy - animal | RRAGC, LAMP1*, RHEB, RRAGD, MLST8 | 1.96E-04 |
| | hsa04145:Phagosome | LAMP1*, ATP6V1D, ATP6V1C1, ATP6V1F | 0.002 |
| | hsa04721:Synaptic vesicle cycle | ATP6V1D, ATP6V1C1, ATP6V1F | 0.008 |
| | hsa00190:Oxidative phosphorylation | ATP6V1D, ATP6V1C1, ATP6V1F | 0.025 |
| **Module 2** | hsa03010:Ribosome | RPS26, RPS7, RPS21, RPL28, RPL26L1 | 5.26E-04 |
| | hsa04514:Cell adhesion molecules | HLA-C, NCAM1, TIGIT | 0.048 |
| | hsa04145:Phagosome | SEC61G, HLA-C, SEC61B | 0.048 |
| **Module 3** | hsa00480:Glutathione metabolism | G6PD, GPX4, GSTP1 | 0.001 |
| | hsa01100:Metabolic pathways | NQO1, G6PD, GPX4, GSTP1, AKR1B1, ENO3 | 0.005 |
| | hsa04216:Ferroptosis | GPX4, FTH1 | 0.037 |
| | **Fusion-Positive Vs Normal: Downregulated DEGs** | | |
| **Module 1** | hsa04530:Tight junction | OCLN*, MARVELD2, F11R, TJP3* | 2.76E-05 |
| **Module 2** | hsa00280:Valine, leucine and isoleucine degradation | AUH, IVD, BCKDHB, ACAT1 | 1.49E-06 |
| | hsa00650:Butanoate metabolism | BDH1, ACAT1 | 0.015 |
| | hsa01100:Metabolic pathways | AUH, BDH1, IVD, BCKDHB, ACAT1 | 0.004 |
| | hsa00071:Fatty acid degradation | CPT2, ACAT1 | 0.024 |
| | hsa01212:Fatty acid metabolism | CPT2, ACAT1 | 0.031 |
| **Module 3** | hsa04071:Sphingolipid signaling pathway | MAPK10, MAPK11, RAC3, GNAI1 | 2.57E-06 |
| | hsa04010:MAPK signaling pathway | MAPK10, MAPK11, RAC3 | 0.003 |
| | hsa05200:Pathways in cancer | MAPK10, RAC3*, GNAI1 | 0.01 |
| | hsa04370:VEGF signaling pathway | MAPK11, RAC3 | 0.02 |
| | hsa04658:Th1 and Th2 cell differentiation | MAPK10, MAPK11 | 0.03 |
| | hsa04657:IL-17 signaling pathway | MAPK10, MAPK11 | 0.031 |
| | hsa04659:Th17 cell differentiation | MAPK10, MAPK11 | 0.036 |
| | hsa04668:TNF signaling pathway | MAPK10, MAPK11 | 0.039 |
| | hsa04660:T cell receptor signaling pathway | MAPK10, MAPK11 | 0.04 |
| | **Fusion-Negative Vs Normal: Upregulated DEGs** | | |
| **Module 1** | hsa04110:Cell cycle | CCNB2*, CDC45*, MCM7, CDCA5, BUB1B*, TTK, CDC7, TRIP13, NDC80, AURKB* | 2.10E-11 |
| | hsa03460:Fanconi anemia pathway | FANCI, RAD51*, UBE2T, BRCA1* | 2.41E-04 |
| **Module 2** | hsa04814:Motor proteins | DNAH1, DNAI1 | 0.022285068 |
| | hsa05014:Amyotrophic lateral sclerosis | DNAH1, DNAI1 | 0.041968326 |
| **Module 3** | hsa04060:Cytokine-cytokine receptor interaction | IL1A, CD40, CXCL8*, IL4R, IL10RA, IL18, TNFSF10, CCL3 | 3.57E-06 |
| | hsa05144:Malaria | CD40, CXCL8*, IL18, ITGAL, TLR2* | 5.05E-06 |
| | hsa05417:Lipid and atherosclerosis | CD40, CXCL8*, IL18, TNFSF10, CCL3, AGER, TLR2* | 7.93E-06 |
| | hsa04640:Hematopoietic cell lineage | IL1A, ITGAM*, IL4R, CD1D, CD33 | 7.65E-05 |
| | **Fusion-Negative Vs Normal: Downregulated DEGs** | | |
| **Module 1** | hsa04014:Ras signaling pathway | NTRK1, PDGFRB, NTRK2, NGFR, FGF7, ANGPT1, HGF, IGF2, FGF1 | 8.61E-07 |
| | hsa04015:Rap1 signaling pathway | PDGFRB, NGFR, FGF7, ANGPT1, CDH1*, HGF, FGF1, THBS1 | 5.26E-06 |
| | hsa05200:Pathways in cancer | NTRK1, PDGFRB, TGFB2, FGF7, CDH1, HGF, IGF2, FOS, FGF1 | 2.97E-04 |
| | hsa04151:PI3K-Akt signaling pathway | NTRK1, PDGFRB, NTRK2, NGFR, FGF7, ANGPT1, IRS1, HGF, IGF2, FGF1, THBS1 | 1.69E-07 |
| | hsa03320:PPAR signaling pathway | ACOX2, EHHADH, SLC27A2 | 0.027165488 |
| | hsa01100:Metabolic pathways | DAO, ACOX2, EPHX2, EHHADH, CAT, ACOT2, PIPOX, AGXT, HAO2, ACOT4, DHRS4 | 0.028408286 |

| | | | |
|---|---|---|---|
| **Module 2** | hsa05200:Pathways in cancer | PDGFRA, NOTCH3, IL6, NOTCH1*, GADD45B, FGF18, F2R, PTGS2, ETS1, ESR1, VEGFA | 3.98E-04 |
| | hsa05224:Breast cancer | NOTCH3, NOTCH1, GADD45B, FGF18, PGR, ESR1 | 0.001070507 |
| | hsa04514:Cell adhesion molecules | SELP, CLDN5, CDH5, SELE, JAM2, JAM3 | 0.001393289 |
| | hsa04933:AGE-RAGE signaling pathway in diabetic complications | IL6*, SERPINE1, NOX4, SELE, VEGFA* | 0.001964469 |
| | hsa04020:Calcium signaling pathway | PDGFRA, FGF18, F2R, NOS1, NGF, MYLK, VEGFA | 0.002120 048 |
| **Module 3** | hsa04820:Cytoskeleton in muscle cells | SPTBN4, LAMA2, ITGA1, FBLN1, ANK2, FBLN2, ITGA10, COL5A3, ITGA11, ITGA8, COL4A5, MYH11, MYH10, ITGA9 | 3.01E-10 |
| | hsa04512:ECM-receptor interaction | VTN, LAMA2, ITGA10, ITGA11, ITGA1, ITGA8, COL4A5, CD36, ITGA9 | 2.92E-08 |
| | hsa04510:Focal adhesion | VTN, LAMA2, ITGA10, FLT4, ITGA11, ITGA1, ITGA8, COL4A5, ITGA9 | 1.58E-05 |
| | hsa05412:Arrhythmogenic right ventricular cardiomyopathy | LAMA2, ITGA10, ITGA11, ITGA1, ITGA8, ITGA9 | 1.16E-04 |
| | hsa05410:Hypertrophic cardiomyopathy | LAMA2, ITGA10, ITGA11, ITGA1, ITGA8, ITGA9 | 2.26E-04 |

**Supplementary Table S1B.** List of identified significant GO terms of the genes of modules in each dataset.

| Enriched GO terms of upregulated DEGs of module in fusion-positive KIRP samples | | | | |
|---|---|---|---|---|
| **Modules** | **Category** | **Term** | **Genes** | **PValue** |
| **Module1** | BP | GO:0032008~positive regulation of TOR signaling | FLCN, RRAGC, RHEB, RRAGD, MLST8, LAMTOR2 | 9.93E-11 |
| | BP | GO:1904263~positive regulation of TORC1 signaling | FLCN, RRAGC, RHEB, RRAGD, LAMTOR2, FNIP2 | 6.54E-10 |
| | BP | GO:0007165~signal transduction | CD274, RHEB, IL2RB, CD38 | 0.075 |
| | BP | GO:0097401~synaptic vesicle lumen acidification | ATP6V1D, ATP6V1C1, ATP6V1F | 8.32E-05 |
| | BP | GO:0007042~lysosomal lumen acidification | LAMP1, ATP6V1D, ATP6V1F | 1.68E-04 |
| | BP | GO:0031669~cellular response to nutrient levels | RRAGC, RHEB, MLST8 | 3.41E-04 |
| | BP | GO:0016241~regulation of macroautophagy | RHEB, ATP6V1D, ATP6V1C1 | 9.58E-04 |
| | CC | GO:0005829~cytosol | PRF1, FLCN, LAMP1, RRAGC, CASTOR2, RHEB, RRAGD, IL2RB, MLST8, FNIP2, ATP6V1D, ATP6V1C1, ATP6V1F | 8.81E-05 |
| | CC | GO:0005886~plasma membrane | CD274, FLCN, LAMP1, RHEB, CD8A, IL2RB, PRF1, CD38, LAMTOR2, ATP6V1D, ATP6V1C1, ATP6V1F | 5.43E-04 |
| | CC | GO:0005765~lysosomal membrane | FLCN, RRAGC, LAMP1, RHEB, RRAGD, MLST8, LAMTOR2, FNIP2, ATP6V1D, ATP6V1C1, ATP6V1F | 4.62E-14 |
| | CC | GO:0016020~membrane | RRAGC, LAMP1, RHEB, IL2RB, PRF1, CD38, ATP6V1D, ATP6V1F | 0.067 |
| | CC | GO:0070062~extracellular exosome | CD274, LAMP1, RHEB, CD38, ATP6V1D, ATP6V1C1, ATP6V1F | 0.004 |
| | CC | GO:1990877~FNIP-folliculin RagC/D GAP | FLCN, RRAGC, LAMTOR2, FNIP2 | 4.41E-08 |
| | CC | GO:0010008~endosome membrane | LAMP1, LAMTOR2, ATP6V1D, ATP6V1F | 0.001 |
| | MF | GO:0005515~protein binding | CD274, PRF1, FLCN, LAMP1, RRAGC, CASTOR2, RHEB, CD8A, RRAGD, IL2RB, MLST8, LAMTOR2, FNIP2, ATP6V1D, ATP6V1C1, ATP6V1F | 0.012 |
| | MF | GO:0046961~proton-transporting ATPase | ATP6V1D, ATP6V1C1, ATP6V1F | 1.91E-04 |
| | MF | GO:0019003~GDP binding | RRAGC, RHEB, RRAGD | 0.001 |
| | MF | GO:0060090~molecular adaptor activity | RRAGC, RRAGD, LAMTOR2 | 0.010 |
| | MF | GO:0003924~GTPase activity | RRAGC, RHEB, RRAGD | 0.034 |
| | MF | GO:0005525~GTP binding | RRAGC, RHEB, RRAGD | 0.043 |
| | MF | GO:0051020~GTPase binding | RRAGC, RRAGD | 0.032 |
| **Module2** | BP | GO:0002181~cytoplasmic translation | RPS26, RPS7, RACK1, RPS21, RPL28, RPL26L1 | 1.92E-07 |
| | BP | GO:0006412~translation | RPS26, RPS7, NACA, RPS21, RPL28 | 1.55E-04 |
| | BP | GO:0006954~inflammatory response | CXCL10, CSF1, NKG7, IDO1 | 0.019 |
| | BP | GO:0042274~ribosomal small subunit biogenesis | NOP56, UTP4, RPS7, AATF | 1.50E-04 |
| | BP | GO:0006364~rRNA processing | NOP56, RPS7, RRP1 | 0.012 |
| | BP | GO:0042267~natural killer cell mediated cytotoxicity | LAG3, NKG7 | 0.045 |
| | BP | GO:0072344~rescue of stalled ribosome | RACK1, PELO | 0.044 |
| | BP | GO:0008015~blood circulation | CXCL10, STAT1 | 0.042 |
| | BP | GO:0000462~maturation of SSU-rRNA | UTP4, AATF | 0.037 |
| | BP | GO:0000027~ribosomal large subunit assembly | BOP1, RRS1 | 0.024 |
| | CC | GO:0016020~membrane | NOP56, RPS26, RPS7, CSF1, SEC61G, NKG7, HLA-C, NCAM1, SEC61B, TIGIT, B2M, RPL28 | 0.042 |
| | CC | GO:0005654~nucleoplasm | NOP56, RPS26, BOP1, UTP4, RPS7, STAT1, PINX1, RRS1, RACK1, AATF, RPS21 | 0.024 |
| | CC | GO:0005730~nucleolus | NOP56, BOP1, UTP4, RPS7, STAT1, PINX1, RRP1, RRS1, AATF | 2.80E-04 |
| | CC | GO:0009897~external side of plasma membrane | CXCL10, LAG3, HLA-C, NCAM1, B2M | 0.002 |
| | CC | GO:0045202~synapse | RPS26, RPS7, RPS21, RPL28 | 0.026 |

| | | | | |
|---|---|---|---|---|
| | CC | GO:0022626~cytosolic ribosome | RPS7, RPS21, RPL28, PELO | 2.39E-04 |
| | CC | GO:0032040~small-subunit processome | NOP56, UTP4, RPS7, AATF | 1.32E-04 |
| | MF | GO:0005515~protein binding | CSF1, PINX1, RRP1, AATF, SEC61G, RACK1, RRS1, NCAM1, SEC61B, TIGIT, PELO, B2M, NOP56, LAG3, UTP4, RPS7, STAT1, GZMA, NKG7, HLA-C, RPS26, BOP1, CXCL10, NACA, RPL28, RPS21, RPL26L1 | 2.04E-04 |
| | MF | GO:0003723~RNA binding | NOP56, UTP4, RPS7, RRP1, AATF, RPS26, BOP1, RRS1, RACK1, SEC61B, RPL28, RPS21, RPL26L1 | 1.92E-07 |
| | MF | GO:0042803~protein homodimerization activity | CSF1, STAT1, GZMA, RACK1, B2M | 0.019 |
| | MF | GO:0003735~structural constituent of ribosome | RPS26, RPS7, RPS21, RPL28, RPL26L1 | 1.36E-04 |
| | MF | GO:0005102~signaling receptor binding | CXCL10, RACK1, HLA-C, TIGIT | 0.014 |
| | MF | GO:0045296~cadherin binding | NOP56, RPS26, STAT1, RACK1 | 0.011 |
| | MF | GO:0043022~ribosome binding | SEC61G, RACK1, SEC61B, PELO | 1.60E-04 |
| **Module3** | BP | GO:0043066~negative regulation of apoptosis | NQO1, GSTP1, AKR1B1, SOD2 | 0.003 |
| | BP | GO:0034599~cellular response to oxidative stress | NQO1, G6PD, GPX4, SOD2 | 3.02E-05 |
| | BP | GO:0006979~response to oxidative stress | NQO1, PRDX4, GPX4 | 0.002 |
| | BP | GO:0098869~cellular oxidant detoxification | PRDX4, GPX4, GSTP1 | 8.94E-04 |
| | BP | GO:0021762~substantia nigra development | G6PD, MBP, ENO3 | 3.76E-04 |
| | BP | GO:0006749~glutathione metabolic process | G6PD, GSTP1, SOD2 | 3.02E-04 |
| | BP | GO:0048147~negative regulation of fibroblast | GSTP1, FTH1, SOD2 | 2.23E-04 |
| | BP | GO:0110076~negative regulation of ferroptosis | NQO1, GPX4, FTH1 | 9.47E-06 |
| | BP | GO:0009636~response to toxic substance | NQO1, MBP | 0.046 |
| | BP | GO:0043525~positive regulation of neuron apoptosis | NQO1, CDK5 | 0.038 |
| | CC | GO:0005829~cytosol | NQO1, G6PD, PRDX4, CDK5, GPX4, MAP2, GSTP1, FTH1, AKR1B1, MBP, TPPP, ENO3 | 5.08E-06 |
| | CC | GO:0005634~nucleus | NQO1, PRDX4, CDK5, GPX4, GSTP1, FTH1, MBP, TPPP | 0.035 |
| | CC | GO:0070062~extracellular exosome | G6PD, PRDX4, GPX4, GSTP1, FTH1, AKR1B1, SOD2, ENO3 | 8.27E-05 |
| | CC | GO:0005739~mitochondrion | GPX4, GSTP1, TPPP, SOD2 | 0.048 |
| | CC | GO:0043025~neuronal cell body | NQO1, CDK5, MAP2, MBP | 0.001 |
| | CC | GO:0030425~dendrite | NQO1, CDK5, MAP2 | 0.027 |
| | CC | GO:0005874~microtubule | CDK5, MAP2, TPPP | 0.014 |
| | MF | GO:0005515~protein binding | NQO1, G6PD, PRDX4, CDK5, GPX4, MAP2, GSTP1, FTH1, AKR1B1, MBP, TPPP, SOD2 | 0.049 |
| | MF | GO:0042802~identical protein binding | NQO1, G6PD, PRDX4, GPX4, FTH1, SOD2 | 0.002 |
| | MF | GO:0048156~tau protein binding | CDK5, MAP2 | 0.026 |
| | MF | GO:0004602~glutathione peroxidase activity | GPX4, GSTP1 | 0.014 |
| | MF | GO:0004784~superoxide dismutase activity | NQO1, SOD2 | 0.003 |
| **Enriched GO terms of downregulated DEGs of module in fusion-positive KIRP samples** | | | | |
| **Modules** | **Category** | **Term** | **Genes** | **PValue** |
| **Module1** | BP | GO:0070830~bicellular tight junction assembly | OCLN, MARVELD2 | 0.010 |
| | BP | GO:0090557~establishment of endothelial barrier | F11R, TJP3 | 0.002 |
| | BP | GO:0150105~protein localization to cell-cell junction | DSP, TJP3 | 0.001 |
|  | BP | GO:1905605~regulation of barrier permeability | OCLN, TJP3 | 0.001 |
| | BP | GO:0098609~cell-cell adhesion | DSP, F11R, TJP3 | 6.99E-04 |
| | BP | GO:0035633~maintenance of blood-brain barrier | OCLN, F11R, TJP3 | 1.73E-05 |
| | BP | GO:0045216~cell-cell junction organization | OCLN, MARVELD2, TJP3 | 1.34E-05 |

| | | | | |
|---|---|---|---|---|
| | CC | GO:0005911~cell-cell junction | OCLN, F11R | 0.036 |
| | CC | GO:0070160~tight junction | OCLN, MARVELD2, F11R | 1.04E-05 |
| | CC | GO:0030054~cell junction | OCLN, MARVELD2, F11R, TJP3 | 2.81E-06 |
| | CC | GO:0005923~bicellular tight junction | OCLN, MARVELD2, F11R, TJP3 | 1.03E-06 |
| **Module2** | BP | GO:0009083~ amino acid catabolic process | IVD, BCKDHB | 0.004 |
| | BP | GO:0006552~L-leucine catabolic process | AUH, IVD | 0.002 |
| | BP | GO:0006635~fatty acid beta-oxidation | BDH2, CPT2, AUH, ECHDC2, ACAT1 | 1.07E-09 |
| | CC | GO:0005759~mitochondrial matrix | AUH, BDH1, IVD, BCKDHB, ACAT1 | 5.43E-06 |
| | CC | GO:0005739~mitochondrion | CPT2, AUH, BDH1, IVD, BCKDHB, ECHDC2, ACAT1 | 8.41E-07 |
| | MF | GO:0003824~catalytic activity | BCKDHB, ECHDC2 | 0.016 |
| | MF | GO:0003858~3-hydroxybutyrate activity | BDH2, BDH1 | 0.001 |
| **Enriched GO terms of upregulated DEGs of module in fusion-negative KIRP samples** | | | | |
| **Modules** | **Category** | **Term** | **Genes** | **PValue** |
| **Module1** | BP | GO:0051301~cell division | CDCA3, CDCA5, KIF14, CDCA8, NCAPG, BUB1B, CDC7, PIMREG, NDC80, NCAPH, AURKB, CCNB2, KIF18B, NUF2, KNTC1, NEK2, KIF2C, SPC25 | 1.43E-17 |
| | BP | GO:0007059~chromosome segregation | CENPU, CENPH, NUF2, CENPK, CENPM, TTK, KIF2C, NEK2, BRCA1, NDC80, DLGAP5, SPC25 | 9.58E-16 |
| | BP | GO:0006974~DNA damage response | RAD51AP1, POLQ, RAD51, MCM7, UHRF1, UBE2T, PCLAF, MCM10, BRCA1 | 6.88E-07 |
| | BP | GO:0007049~cell cycle | FANCI, MCM7, CDCA3, UHRF1, NUF2, BRCA1, PIMREG, CDKN3 | 1.25E-05 |
| | BP | GO:0016310~phosphorylation | PBK, BUB1B, TTK, CDC7, NEK2, AURKB, CDKN3 | 0.004 |
| | BP | GO:0006281~DNA repair | RAD51AP1, FANCI, POLQ, RAD51, EXO1, UBE2T, BRCA1 | 4.42E-05 |
| | BP | GO:0000724~double-strand break repair | RAD51AP1, RAD51, UHRF1, BRCA1 | 0.002 |
| | BP | GO:0006302~double-strand break repair | POLQ, CDCA5, BRCA1, TRIP13 | 8.71E-04 |
| | BP | GO:0051321~meiotic cell cycle | RAD51AP1, RAD51, EXO1, NEK2 | 8.38E-04 |
| | BP | GO:0036297~interstrand cross-link repair | RAD51AP1, FANCI, NEIL3, RAD51 | 1.07E-04 |
| | CC | GO:0005634~nucleus | ARHGAP11A, MCM7, UHRF1, CDCA5, KIF14, CDCA7, NCAPG, CDCA8, BUB1B, MCM10, TTK, BRCA1, TYMS, CENPA, AURKB, RAD51AP1, CCNB2, CDC45, EXO1, NUF2, PBK, PCLAF, KNTC1, NEK2, DLGAP5, CENPU, GINS2, POLQ, CDC7, NDC80, NEIL3, RAD51, KIF18B, CENPH, UBE2T, CENPK, DEPDC1, CENPM, KIF2C, TRIP13, CDKN3, SPC25 | 1.28E-15 |
| | CC | GO:0005829~cytosol | ARHGAP11A, CDCA3, MCM7, CDCA5, KIF14, CDCA7, NCAPG, CDCA8, BUB1B, HMMR, TYMS, CENPA, NCAPH, AURKB, CCNB2, NUF2, KNTC1, NEK2, DLGAP5, FANCI, CENPU, POLQ, NDC80, RAD51, KIF18B, CENPH, CENPK, CENPM, KIF2C, CDKN3, SPC25 | 2.82E-07 |
| | CC | GO:0005654~nucleoplasm | MCM7, UHRF1, CDCA5, CDCA7, CDCA8, MCM10, BRCA1, CENPA, NCAPH, AURKB, RAD51AP1, CDC45, EXO1, NUF2, PCLAF, NEK2, E2F7, FANCI, CENPU, GINS2, POLQ, CDC7, PIMREG, NDC80, NEIL3, RAD51, KIF18B, CENPH, UBE2T, CENPK, CENPM | 6.13E-11 |
| | CC | GO:0005737~cytoplasm | FANCI, TROAP, CDCA5, CDCA7, KIF14, NCAPG, BUB1B, TTK, CDC7, BRCA1, HMMR, TYMS, NDC80, NCAPH, CCNB2, RAD51, KIF18B, KNTC1, NEK2, KIF2C, DLGAP5, CDKN3 | 0.011 |
| | CC | GO:0005813~centrosome | CCNB2, RAD51, CDC45, PCLAF, BUB1B, KIF2C, NEK2, HMMR, NDC80 | 1.32E-04 |
| | CC | GO:0000776~kinetochore | CENPH, NUF2, BUB1B, TTK, KIF2C, NEK2, NDC80, AURKB, SPC25 | 4.06E-09 |
| | CC | GO:0005694~chromosome | RAD51AP1, CENPU, NEIL3, MCM7, CENPH, CDCA5, BRCA1, TRIP13 | 1.74E-06 |
| | MF | GO:0005515~protein binding | CDCA3, MCM7, UHRF1, TROAP, CDCA5, KIF14, CDCA7, NCAPG, CDCA8, BUB1B, MCM10, TTK, HMMR, BRCA1, CENPA, | 8.53E-06 |

| | | | | |
|---|---|---|---|---|
| | | | NCAPH, AURKB, RAD51AP1, CCNB2, CDC45, EXO1, NUF2, PBK, PCLAF, KNTC1, NEK2, E2F7, DLGAP5, FANCI, CENPU, GINS2, POLQ, CDC7, PIMREG, NDC80, RAD51, KIF18B, CENPH, UBE2T, CENPK, DEPDC1, KIF2C, TRIP13, CDKN3, SPC25 | |
| | MF | GO:0005524~ATP binding | POLQ, MCM7, KIF14, BUB1B, TTK, CDC7, AURKB, RAD51, KIF18B, UBE2T, PBK, NEK2, KIF2C, TRIP13 | 5.44E-05 |
| | MF | GO:0003682~chromatin binding | POLQ, RAD51, CDC45, EXO1, UBE2T, CDCA5, PCLAF, CENPA, NCAPH | 3.41E-05 |
| | MF | GO:0003677~DNA binding | RAD51AP1, FANCI, POLQ, MCM7, EXO1, UHRF1, BRCA1, CENPA | 0.020 |
| | MF | GO:0016887~ATP hydrolysis activity | POLQ, RAD51, KIF18B, MCM7, KIF14, KIF2C, TRIP13 | 6.14E-04 |
| | MF | GO:0004674~protein serine/threonine kinase activity | PBK, BUB1B, TTK, CDC7, NEK2, AURKB | 0.002 |
| | MF | GO:0106310~protein serine kinase activity | PBK, BUB1B, TTK, CDC7, NEK2, AURKB | 0.001 |
| **Module2** | BP | GO:0003341~cilium movement | DNAAF1, HYDIN, ZMYND10, DNAI1, CCDC40 | 3.23E-09 |
| | BP | GO:0070286~axonemal dynein assembly | DNAAF3, DNAAF1, CCDC65, DRC1, CCDC40 | 2.72E-11 |
| | BP | GO:0044458~motile cilium assembly | DNAAF3, DNAAF1, ZMYND10, CCDC40 | 2.14E-07 |
| | BP | GO:0036159~inner dynein arm assembly | DNAH1, DNAAF1, ZMYND10, CCDC40 | 4.45E-08 |
| | BP | GO:0060271~cilium assembly | DNAAF1, CCDC65, CCNO | 0.005 |
| | BP | GO:0007507~heart development | DNAAF3, DRC1, DNAI1 | 0.003 |
| | BP | GO:0030317~flagellated sperm motility | DNAH1, DNAI1, CCDC40 | 8.59E-04 |
| | BP | GO:0007368~determination of left/right symmetry | DNAAF3, DRC1, DNAI1 | 4.15E-04 |
| | BP | GO:0036158~outer dynein arm assembly | DNAAF1, ZMYND10, DNAI1 | 6.45E-05 |
| | BP | GO:0003351~epithelial cilium movement | DNAH1, DNAI1, CCDC40 | 3.87E-05 |
| | CC | GO:0005930~axoneme | DNAH1, DNAAF1, HYDIN, CCDC65, DRC1, CCDC40 | 1.07E-09 |
| | CC | GO:0005929~cilium | DNAAF1, DNAI1, CCDC40 | 0.004 |
| | CC | GO:0120293~dynein axonemal particle | DNAAF3, ZMYND10, DNAI1 | 3.12E-05 |
| | CC | GO:0005858~axonemal dynein complex | DNAH1, CCDC65, DRC1 | 2.24E-05 |
| **Module3** | BP | GO:0006954~inflammatory response | IL1A, CD40, CXCL8, IL18, CCL3, SIGLEC1, ITGAL, AGER, HAVCR2, TLR2 | 5.38E-10 |
| | BP | GO:0006955~immune response | IL1A, ZAP70, CXCL8, IL4R, IL18, TNFSF10, CCL3, CD1D, TLR2 | 6.08E-08 |
| | BP | GO:0045087~innate immune response | ZAP70, ITGAM, CD1D, CORO1A, CD244, CAMP, HAVCR2, TLR2 | 1.76E-06 |
| | BP | GO:0007165~signal transduction | CXCL8, IL4R, TNFSF10, ITGAL, CD244, CD33, TLR2 | 0.002 |
| | BP | GO:0071222~cellular response to lipopolysaccharide | IL1A, CD40, CXCL8, IL18, CAMP, HAVCR2 | 2.35E-06 |
| | BP | GO:0032760~ regulation of tumor necrosis factor | IL1A, CCL3, AGER, HAVCR2, TLR2 | 8.01E-06 |
| | BP | GO:0032722~positive regulation of chemokine production | IL4R, IL18, AGER, HAVCR2, TLR2 | 1.68E-07 |
| | BP | GO:0007155~cell adhesion | ITGAM, ITGAL, AGER, CD33 | 0.021 |
| | BP | GO:0010628~positive regulation of gene expression | IL1A, CXCL8, CCL3, TLR2 | 0.020 |
| | BP | GO:0008285~negative regulation of cell population | IL1A, CXCL8, CD33, TLR2 | 0.013 |
| | CC | GO:0005886~plasma membrane | CD52, CD40, ITGAM, IL4R, IL10RA, CD1D, ITGAL, CORO1A, AGER, ZAP70, CD207, TNFSF10, SIGLEC1, HCST, CD244, CD33, MYO1G, TLR2 | 3.25E-06 |
| | CC | GO:0016020~membrane | CD52, ITGAM, IL4R, IL10RA, CD207, TNFSF10, SIGLEC1, ITGAL, CORO1A, MYO1G, HAVCR2, TLR2 | 0.011 |
| | CC | GO:0005576~extracellular region | IL1A, CD52, CXCL8, IL4R, IL18, TNFSF10, CCL3, SIGLEC1, AGER, CAMP | 2.28E-04 |
| | CC | GO:0009986~cell surface | IL1A, CD40, ITGAM, CD1D, ITGAL, AGER, HCST, CD33, HAVCR2, TLR2 | 1.57E-08 |
| | CC | GO:0005615~extracellular space | IL1A, CD40, ITGAM, CXCL8, IL18, TNFSF10, CCL3, CD1D, CAMP | 0.001 |
| | CC | GO:0009897~external side of plasma membrane | CD40, ITGAM, IL4R, CD207, CD1D, ITGAL, CD244, CD33 | 2.71E-07 |
| | CC | GO:0070062~extracellular exosome | CD40, ITGAM, TNFSF10, ITGAL, CORO1A, CAMP, MYO1G | 0.031 |

| | | | | |
|---|---|---|---|---|
| | MF | GO:0005515~protein binding | CD40, ITGAM, IL4R, CXCL8, IL10RA, IL18, CD1D, ITGAL, CORO1A, AGER, IL1A, ZAP70, CD207, TNFSF10, CCL3, HCST, CD244, CD33, CAMP, HAVCR2, TLR2 | 0.008 |
| | MF | GO:0038023~signaling receptor activity | CD40, IL10RA, CD207, AGER, CD244, CD33, TLR2 | 1.50E-07 |
| | MF | GO:0030246~carbohydrate binding | CD207, SIGLEC1, CD33 | 0.022 |
| | MF | GO:0005125~cytokine activity | IL1A, IL18, TNFSF10 | 0.020 |
| | MF | GO:0001540~amyloid-beta binding | ITGAM, AGER, TLR2 | 0.004 |
| **Enriched GO terms of downregulated DEGs of module in fusion-negative KIRP samples** | | | | |
| **Modules** | **Category** | **Term** | **Genes** | **PValue** |
| **Module1** | BP | GO:0051897~positive regulation of kinase B signal | NTRK1, PDGFRB, NTRK2, TGFB2, ANGPT1, IRS1, HGF, CAT, IGF2, THBS1 | 1.50E-11 |
| | BP | GO:0008284~positive regulation of cell population | PDGFRB, NTRK2, TGFB2, FGF7, IRS1, IGF2, FGF1, THBS1 | 1.35E-05 |
| | BP | GO:0007165~signal transduction | PDGFRB, FGF7, IRS1, PECAM1, CCN2, FGF1 | 0.047 |
| | BP | GO:0010628~positive regulation of gene expression | ACTA2, NTRK2, FGF7, ANGPT1, EPHX2, FGF1 | 0.001 |
| | BP | GO:0070374~positive regulation of ERK1 and ERK2 cascade | NTRK1, PDGFRB, ACTA2, ANGPT1, CCN2, FGF1 | 3.08E-05 |
| | BP | GO:0001934~positive regulation of protein phosphorylation | NTRK1, NTRK2, ANGPT1, HGF, PECAM1, FGF1 | 1.89E-05 |
| | BP | GO:0030154~cell differentiation | NTRK1, NTRK2, FGF7, ANGPT1, FGF1 | 0.023 |
| | BP | GO:0043066~negative regulation of apoptotic process | NTRK1, ANGPT1, HGF, CAT, THBS1 | 0.008 |
| | BP | GO:0001525~angiogenesis | PDGFRB, ANGPT1, CCN2, FGF1, HRG | 7.58E-04 |
| | BP | GO:0009410~response to xenobiotic stimulus | NTRK1, CDH1, CAT, FOS, THBS1 | 7.26E-04 |
| | CC | GO:0005829~cytosol | NGFR, NTRK2, IRS1, EPHX2, PIPOX, FOS, FGF1, DHRS4, ACTA2, DAO, ACOX2, EHHADH, CAT, ACOT2, SNAI1, AGXT, HAO2, SLC27A2, ACOT4 | 4.59E-04 |
| | CC | GO:0005576~extracellular region | NGFR, TGFB2, ANGPT1, HGF, ELN, AFM, IGF2, FGF1, THBS1, DHRS4, FGF7, DAO, CDH1, CAT, CCN2, HRG | 1.12E-07 |
| | CC | GO:0005615~extracellular space | ACTA2, TGFB2, FGF7, ANGPT1, HGF, CAT, PECAM1, IGF2, AFM, CCN2, FGF1, THBS1 | 1.35E-04 |
| | CC | GO:0005782~peroxisomal matrix | DAO, ACOX2, EPHX2, EHHADH, CAT, ACOT2, PIPOX, AGXT, HAO2, ACOT4, DHRS4 | 2.25E-19 |
| | CC | GO:0070062~extracellular exosome | ACTA2, ANGPT1, CDH1, EPHX2, CAT, PECAM1, AFM, HRG, THBS1, SLC27A2 | 0.005 |
| | CC | GO:0005777~peroxisome | DAO, ACOX2, EPHX2, EHHADH, CAT, PIPOX, AGXT, HAO2, ACOT4, DHRS4 | 6.15E-14 |
| | CC | GO:0043231~intracellular membrane-bounded organelle | PDGFRB, ACOX2, IRS1, CAT, SNAI1, CCN2, AGXT | 0.002 |
| | MF | GO:0005515~protein binding | IRS1, ELN, AFM, FGF1, THBS1, FGF7, CDH1, CCN2, HAO2, PDGFRB, NTRK1, NGFR, NTRK2, TGFB2, ANGPT1, HGF, IGF2, PIPOX, FOS, DAO, ACOX2, EHHADH, ACOT2, SNAI1, PECAM1, HRG, AGXT | 0.047 |
| | MF | GO:0042802~identical protein binding | NTRK1, DAO, ANGPT1, CDH1, HGF, CAT, FOS, AGXT, DHRS4 | 0.006 |
| | MF | GO:0042803~protein homodimerization activity | NTRK1, NTRK2, TGFB2, ACOX2, EPHX2, CAT, PECAM1, AGXT, THBS1 | 2.30E-05 |
| | MF | GO:0008201~heparin binding | FGF7, CCN2, FGF1, HRG, THBS1 | 2.53E-04 |
| | MF | GO:0008083~growth factor activity | TGFB2, FGF7, HGF, IGF2, FGF1 | 1.54E-04 |
| | MF | GO:0019899~enzyme binding | PDGFRB, EHHADH, CAT, SLC27A2 | 0.023 |
| | MF | GO:0005102~signaling receptor binding | PDGFRB, TGFB2, HGF, HRG | 0.022 |
| **Module2** | BP | GO:0045944~positive regulation of transcription | NOTCH3, CSF3, EGR2, MYOCD, EGR3, NOTCH1, ETS1, ESR1, VEGFA, NR4A2, NR4A1, IL6, NR4A3, PGR, NOS1 | 4.87E-06 |
| | BP | GO:0010628~positive regulation of gene expression | CLDN5, CDH5, IL6, NOTCH1, ACTC1, FGF18, PGR, EMILIN1, NGF, ETS1, ACTG2, VEGFA | 1.06E-07 |
| | BP | GO:0008285~negative regulation of cell population | CDH5, MYOCD, IL6, BTG2, NOTCH1, DUSP1, F2R, NOX4, PLG, NGF, PTGS2, ETS1 | 2.10E-08 |
| | BP | GO:0006357~regulation of transcription | NR4A2, NR4A1, EGR2, NOTCH1, EGR3, NR4A3, PGR, ETS1, ESR1, VEGFA | 0.029 |
| | BP | GO:0000122~negative regulation of transcription | NR4A2, NOTCH3, MYOCD, ZFP36, BTG2, NOTCH1, NR4A3, ESR1, VEGFA | 0.004 |

| | | | | |
|---|---|---|---|---|
| | BP | GO:0008284~positive regulation of cell population proliferation | CLDN5, PDGFRA, CSF3, MYOCD, IL6, NOTCH1, FGF18, F2R, VEGFA | 6.52E-05 |
| | BP | GO:0045893~positive regulation of DNA-templated transcription | MYOCD, IL6, EGR2, NOTCH1, F2R, NOS1, ETS1, ESR1 | 0.003 |
| | BP | GO:0006954~inflammatory response | SELP, NR4A1, IL6, F2R, NOX4, PTX3, SELE | 0.001 |
| | BP | GO:0032496~response to lipopolysaccharide | SELP, NOTCH1, F2R, REN, NOS1, PTGS2, SELE | 2.14E-06 |
| | BP | GO:0010629~negative regulation of gene expression | CLDN5, NOTCH1, PGR, EMILIN1, ESR1, VEGFA | 0.001 |
| | CC | GO:0005886~plasma membrane | PDGFRA, NOTCH3, NOTCH1, MME, F2R, SERPINE1, LPL, PLG, SELE, ESR1, MYLK, SELP, CDH5, CLDN5, APOH, NOX4, REN, PGR, NOS1, JAM2, ABCG2, JAM3 | 0.034 |
| | CC | GO:0005615~extracellular space | CSF3, SERPINE1, LPL, PLG, HPR, NGF, SELE, ACTG2, VEGFA, SELP, IL6, ACTC1, TTR, APOH, FGF18, ALB, EMILIN1, PTX3, REN, GC, JAM3 | 6.57E-08 |
| | CC | GO:0005576~extracellular region | NOTCH3, CSF3, ITIH2, NOTCH1, F2R, SERPINE1, LPL, PLG, HPR, NGF, VEGFA, IL6, TTR, APOH, FGF18, ALB, EMILIN1, PTX3, REN, GC | 5.94E-07 |
| | CC | GO:0070062~extracellular exosome | BTG2, ITIH2, MME, SERPINE1, PLG, HPR, ACTG2, ACTC1, TTR, APOH, ALB, EMILIN1, GC, PCYOX1 | 0.004 |
| | CC | GO:0009986~cell surface | NOTCH3, CDH5, NOTCH1, MME, APOH, F2R, LPL, PLG, JAM2, VEGFA | 4.28E-05 |
| | CC | GO:0000785~chromatin | NR4A2, NR4A1, MYOCD, EGR2, EGR3, NR4A3, PGR, ETS1, ESR1 | 0.009 |
| | CC | GO:0032991~protein-containing complex | NR4A2, PDGFRA, NOTCH1, ALB, NOS1, PTGS2, ESR1 | 0.008 |
| | MF | GO:0005515~protein binding | NOTCH3, BTG2, ITIH2, NOTCH1, MTTP, SERPINE1, LPL, PLG, PTGS2, ETS1, MYLK, LIPE, CDH5, CNN1, ZFP36, TTR, APOH, PDK4, EMILIN1, NOS1, JAM2, JAM3, PDGFRA, MYOCD, EGR2, GADD45B, MME, DUSP1, F2R, NGF, SELE, ESR1, VEGFA, NR4A2, SELP, CLDN5, NR4A1, IL6, NR4A3, ALB, NOX4, REN, PTX3, PGR, ABCG2, PCYOX1 | 0.003 |
| | MF | GO:0042802~identical protein binding | NOTCH3, NOTCH1, ETS1, ESR1, VEGFA, CLDN5, NR4A1, TTR, APOH, ALB, EMILIN1, PTX3, PGR, ABCG2 | 8.02E-04 |
| | MF | GO:0003677~DNA binding | NR4A2, NR4A1, ZFP36, EGR2, NR4A3, ALB, PGR, ETS1, ESR1 | 0.014 |
| | MF | GO:0001228~DNA-binding transcription activity | NR4A2, NR4A1, EGR2, NOTCH1, EGR3, NR4A3, PGR, ETS1, ESR1 | 6.34E-05 |
| | MF | GO:0042803~protein homodimerization activity | PDGFRA, NR4A3, MME, LPL, PTGS2, ABCG2, JAM3, VEGFA | 0.004 |
| | MF | GO:0019899~enzyme binding | NOTCH3, CSF3, ZFP36, NOTCH1, PGR, PLG, PTGS2, ESR1 | 7.58E-05 |
| | MF | GO:0005102~signaling receptor binding | CDH5, F2R, SERPINE1, LPL, REN, PGR, PLG | 5.58E-04 |
| **Module3** | BP | GO:0010628~positive regulation of gene expression | CYP26B1, WT1, FBLN1, ANK2, CD36, NTS, F3, LDLR | 0.001 |
| | BP | GO:0007605~sensory perception of sound | SPTBN4, WFS1, GJB6, PCDH15, MYO7A, SLC26A4, POU3F4 | 1.49E-05 |
| | BP | GO:0001822~kidney development | CYP26B1, WT1, WFS1, CYP4A22, CYP4A11, ITGA8, PROX1 | 2.75E-06 |
| | BP | GO:0033627~cell adhesion mediated by integrin | VTN, ITGA10, ITGA11, ITGA1, ITGA8, ITGBL1, ITGA9 | 4.14E-09 |
| | BP | GO:0008285~negative regulation of cell population proliferation | CYP27B1, WT1, FRZB, GJB6, ITGA1, PROX1 | 0.014 |
| | BP | GO:0006805~xenobiotic metabolic process | CYP26B1, CYP2B6, CYP4F3, AOX1, UGT1A9 | 6.82E-04 |
| | BP | GO:0007420~brain development | S1PR1, NCAM1, PROX1, POU3F4 | 0.032 |
| | BP | GO:0006629~lipid metabolic process | AOX1, CD36, LPA, LDLR | 0.031 |
| | BP | GO:0035725~sodium ion transmembrane transport | SCN9A, SCN7A, SCN4B, SCN2B | 0.007 |
| | BP | GO:0042632~cholesterol homeostasis | APOM, PLA2G12B, DGAT2, LDLR | 0.005 |
| | CC | GO:0005886~plasma membrane | SPTBN4, SERPINC1, FLT4, TMPRSS4, PCDH15, SCN9A, S1PR1, SCN7A, NCAM1, CD36, LDLR, TMC1, ITGA1, ANK2, F3, SPTB, ITGA10, KCNQ2, ITGA11, ITGA8, ITGBL1, SLC26A4, SCN4B, ITGA9, SCN2B | 0.046 |
| | CC | GO:0016020~membrane | SPTBN4, DGAT2, LAMA2, WFS1, TMPRSS4, CYP4A11, CYP4F3, ITGA1, CYP4A22, PCDH15, FBLN1, ANK2, MYO7A, F3, FBLN2, FRZB, KCNQ2, | 0.027 |

| | | | | |
|---|---|---|---|---|
| | | | ITGA11, COL4A5, NCAM1, CD36, UGT1A9, SLC26A4, LDLR | |
| | CC | GO:0005576~extracellular region | CHGA, PLA2G12B, LAMA2, COL14A1, SERPINC1, FST, FLT4, PON1, ADIPOQ, PCDH15, FBLN1, NTS, FBLN2, VTN, APOM, FRZB, COL5A3, COL4A5, COL21A1, ITGBL1, NCAM1, SOST, LPA | 1.72E-07 |
| | CC | GO:0005615~extracellular space | CHGA, COL14A1, SERPINC1, FST, PON1, TMPRSS4, CYP4A22, ADIPOQ, PCDH15, FBLN1, AFP, F3, VTN, FRZB, COL5A3, COL4A5, COL21A1, SOST, CD36, LPA | 8.40E-06 |
| | CC | GO:0062023~collagen-containing extracellular matrix | LAMA2, COL14A1, SERPINC1, ADIPOQ, FBLN1, F3, FBLN2, VTN, COL5A3, COL4A5, COL21A1, NCAM1, SOST | 4.16E-09 |
| | CC | GO:0005789~endoplasmic reticulum membrane | CYP26B1, CYP2B6, DGAT2, WFS1, CYP4F3, PON1, CYP4A22, CYP4A11, UGT1A9, CYP17A1, PTGS1 | 0.003 |
| | CC | GO:0009897~external side of plasma membrane | TMC1, ITGA10, ITGA11, ITGA1, ITGA8, S1PR1, NCAM1, CD36, F3, LDLR, ITGA9 | 7.86E-07 |
| | MF | GO:0005178~integrin binding | VTN, ITGA10, ITGA11, ITGA1, ITGA8, ITGBL1, FBLN1, ITGA9 | 1.07E-06 |
| | MF | GO:0020037~heme binding | CYP27B1, CYP26B1, CYP2B6, CYP4F3, CYP4A22, CYP4A11, CYP17A1, PTGS1 | 7.95E-07 |
| | MF | GO:0005506~iron ion binding | CYP27B1, CYP26B1, CYP2B6, CYP4F3, CYP4A22, CYP4A11, AOX1, CYP17A1 | 3.99E-07 |
| | MF | GO:0005201~extracellular matrix structural constituent | VTN, LAMA2, COL14A1, COL5A3, ADIPOQ, COL4A5, FBLN1, FBLN2 | 1.17E-07 |
| | MF | GO:0051015~actin filament binding | SPTBN4, GJB6, MYH11, MYO7A, MYH10, SPTB | 0.001 |
| | MF | GO:0005543~phospholipid binding | FABP1, SPTBN4, APOM, PON1, SPTB, F3 | 6.13E-05 |
| | MF | GO:0005518~collagen binding | VTN, COL14A1, ITGA10, COL5A3, ITGA11, ITGA1 | 3.26E-06 |

**Table S2.** List of the identified module's DEGs in fusion-positive KIRP using LIMMA and DESeq2 methods.

| **Upregulated DEGs of modules** | | | | | |
|---|---|---|---|---|---|
| **LIMMA** | | | **DESeq2** | | |
| **Genes** | **Log2Fold Change** | **padj** | **Genes** | **Log2Fold Change** | **padj** |
| SOD2 | 2.135 | 3.02E-08 | SOD2 | 2.064 | 5.15E-05 |
| RRAGC | 1.968 | 4.49E-23 | RRAGC | 1.876 | 7.06E-11 |
| RHEB | 1.893 | 3.32E-24 | RHEB | 1.797 | 5.21E-10 |
| NQO1 | 1.792 | 0.0005 | FNIP2 | 1.712 | 0.0015 |
| RPL26L1 | 1.711 | 5.59E-12 | ATP6V1C1 | 1.618 | 9.38E-06 |
| ATP6V1C1 | 1.674 | 6.97E-09 | ATP6V1F | 1.573 | 6.82E-08 |
| ATP6V1F | 1.600 | 7.04E-15 | RRAGD | 1.526 | 0.0010 |
| FNIP2 | 1.590 | 0.0004 | GPX4 | 1.347 | 0.0003 |
| MBP | 1.483 | 1.83E-09 | RPL26L1 | 1.343 | 2.48E-06 |
| RRAGD | 1.471 | 9.47E-05 | MBP | 1.332 | 1.69E-05 |
| G6PD | 1.392 | 1.68E-09 | NQO1 | 1.305 | 0.0306 |
| GPX4 | 1.324 | 7.38E-06 | RPS7 | 1.291 | 9.90E-07 |
| RPS7 | 1.302 | 1.15E-11 | G6PD | 1.223 | 9.14E-06 |
| STAT1 | 1.257 | 5.95E-05 | STAT1 | 1.168 | 0.0010 |
| LAMP1 | 1.217 | 9.32E-08 | LAMTOR2 | 1.147 | 4.36E-07 |
| LAMTOR2 | 1.203 | 6.71E-11 | LAMP1 | 1.122 | 5.29E-05 |
| RACK1 | 1.087 | 2.59E-11 | RACK1 | 1.062 | 4.00E-06 |
| **Downregulated DEGs of modules** | | | | | |
| MARVELD2 | -2.425 | 2.01E-07 | MARVELD2 | -3.250 | 0.0004 |
| OCLN | -2.049 | 1.25E-05 | OCLN | -3.121 | 0.0026 |
| MAPK11 | -1.417 | 4.69E-06 | MAPK10 | -1.783 | 1.12E-05 |
| CCND1 | -1.305 | 0.0002 | MAPK11 | -1.422 | 0.0076 |
| MAPK10 | -1.212 | 5.49E-05 | CCND1 | -1.334 | 0.0009 |
| F11R | -1.042 | 1.30E-07 | F11R | -1.042 | 4.27E-06 |

**Table S3.** List of key differentially expressed miRNAs (DEMs) targeting the key genes in fusion-positive KIRP modules

| **Upregulated DEMs** | | | | | |
|---|---|---|---|---|---|
| **LIMMA** | | | **DESeq2** | | |
| **miRNA** | **Log2Fold Change** | **padj** | **miRNA** | **Log2Fold Change** | **padj** |
| hsa-mir-5589 | 4.553 | 4.16E-05 | hsa-mir-5589 | 6.557 | 0.0059 |
| hsa-mir-185 | 2.677 | 4.90E-09 | hsa-mir-185 | 2.751 | 7.87488E-18 |
| hsa-mir-221 | 1.767 | 3.98E-05 | hsa-mir-221 | 2.025 | 2.21493E-07 |
| hsa-mir-342 | 1.594 | 0.0102 | hsa-mir-342 | 1.425 | 0.0198 |
| hsa-mir-148a | 1.160 | 0.0011 | hsa-mir-148a | 1.258 | 4.16421E-05 |
| hsa-mir-130b | 1.122 | 0.0048 | hsa-mir-130b | 1.139 | 0.0103 |
| **Downregulated DEMs** | | | | | |
| hsa-mir-130a | -1.406 | 0.0020 | hsa-mir-874 | -1.538 | 0.0057 |
| hsa-mir-26b | -1.374 | 0.0075 | hsa-mir-30c-1 | -1.421 | 0.0131 |
| hsa-mir-30c | -1.536 | 0.0318 | hsa-mir-130a | -1.413 | 0.0002 |
| hsa-mir-30c-2 | -1.465 | 0.0294 | hsa-mir-30c-2 | -1.242 | 0.0142 |
| hsa-mir-874 | -1.605 | 0.0115 | hsa-mir-26b | -1.138 | 0.0067 |

**Supplementary Table S4. Exclusive set of identified significant DEMs in Fusion-Positive and Fusion-Negative KIRP.**

| **Exclusive upregulated DEMs in fusion-positive KIRP** | **Exclusive downregulated DEMs in fusion-positive KIRP** | **Exclusive upregulated DEMs in fusion-negative KIRP** | | **Exclusive downregulated DEMs in fusion-negative KIRP** | | |
|---|---|---|---|---|---|---|
| hsa-mir-466 | hsa-mir-136 | hsa-mir-4677 | hsa-mir-1295a | hsa-mir-934 | hsa-mir-195 | hsa-mir-337 |
| hsa-mir-5589 | hsa-mir-200b | hsa-mir-651 | hsa-mir-223 | hsa-mir-154 | hsa-mir-654 | hsa-mir-509-1 |
| hsa-mir-488 | hsa-mir-874 | hsa-mir-25 | hsa-mir-93 | hsa-mir-362 | hsa-mir-412 | hsa-mir-432 |
| hsa-mir-185 | hsa-mir-497 | hsa-mir-106b | hsa-mir-6892 | hsa-mir-199a-2 | hsa-mir-299 | hsa-mir-4662a |
| hsa-mir-144 | hsa-mir-195 | hsa-mir-4668 | hsa-mir-219a-1 | hsa-mir-10b | hsa-mir-424 | hsa-mir-215 |
| hsa-mir-222 | hsa-mir-27b | hsa-mir-135b | hsa-mir-146a | hsa-mir-539 | hsa-mir-203b | hsa-mir-29c |
| hsa-mir-486-1 | hsa-mir-1468 | hsa-mir-3127 | hsa-mir-96 | hsa-mir-1251 | hsa-mir-203a | hsa-mir-370 |
| hsa-mir-451a | hsa-mir-10a | hsa-mir-4772 | hsa-mir-34b | hsa-mir-199a-1 | hsa-mir-758 | hsa-mir-493 |
| hsa-mir-486-2 | hsa-mir-200a | hsa-mir-181d | hsa-mir-155 | hsa-mir-127 | hsa-mir-134 | hsa-mir-381 |
| hsa-mir-221 | hsa-mir-30c-1 | hsa-mir-589 | hsa-mir-6509 | hsa-mir-411 | hsa-mir-410 | hsa-mir-495 |
| hsa-mir-3200 | hsa-mir-130a | hsa-mir-642a | hsa-mir-935 | hsa-mir-376c | hsa-mir-509-3 | hsa-mir-3065 |
| hsa-mir-3074 | hsa-mir-30c-2 | hsa-mir-1180 | hsa-mir-183 | hsa-mir-136 | hsa-mir-874 | hsa-mir-382 |
| hsa-mir-342 | hsa-mir-26b | hsa-mir-885 | hsa-mir-31 | hsa-mir-502 | hsa-mir-30b | hsa-mir-577 |
| hsa-mir-148a | | hsa-mir-1293 | hsa-mir-211 | hsa-mir-190a | hsa-mir-450a-1 | hsa-mir-135a-2 |
| hsa-mir-130b | | hsa-mir-599 | hsa-mir-1258 | hsa-mir-1296 | hsa-mir-450a-2 | hsa-mir-500b |
| | | hsa-mir-7156 | hsa-mir-561 | hsa-mir-26b | hsa-mir-660 | hsa-mir-655 |
| | | hsa-mir-875 | hsa-mir-6718 | | | |

### Supplementary Table S5. PRCC and TFE3 targeting and expressed miRNAs in Fusion-Positive KIRP.

| **PRCC targeting expressed miRNAs (total = 53)** | | **TFE3 targeting expressed miRNAs (total = 122 )** | | | | |
|---|---|---|---|---|---|---|
| hsa-miR-1228-3p | hsa-miR-362-3p | hsa-miR-30c-2-3p | hsa-miR-675-3p | hsa-miR-659-3p | hsa-miR-5010-5p | hsa-miR-500b-5p |
| hsa-miR-125a-3p | hsa-miR-365b-5p | hsa-miR-30c-2-3p | hsa-miR-874-3p | hsa-miR-758-3p | hsa-miR-6716-5p | hsa-miR-4728-5p |
| hsa-miR-125b-2-3p | hsa-miR-378a-3p | hsa-miR-221-3p | hsa-miR-1228-3p | hsa-miR-1288-3p | hsa-miR-6842-5p | hsa-miR-184 |
| hsa-miR-1296-3p | hsa-miR-421- | hsa-miR-185-3p | hsa-miR-1266-3p | hsa-miR-500b-3p | hsa-miR-937-5p | hsa-miR-429 |
| hsa-miR-150-5p | hsa-miR-452-5p | hsa-let-7b-3p | hsa-miR-3117-3p | hsa-miR-5010-3p | hsa-miR-24-2-5p | hsa-miR-484 |
| hsa-miR-151a-3p | hsa-miR-4661-3p | hsa-let-7f-1-3p | hsa-miR-3605-3p | hsa-miR-135b-5p | hsa-miR-25-5p | hsa-miR-4636 |
| hsa-miR-15b-5p | hsa-miR-497-5p | hsa-miR-129-1-3p | hsa-miR-3677-3p | hsa-miR-23a-5p | hsa-miR-92a-2-5p | hsa-miR-3170 |
| hsa-miR-181c-5p | hsa-miR-508-5p | hsa-miR-125b-1-3p | hsa-miR-3922-3p | hsa-miR-103a-2-5p | hsa-miR-197-5p | hsa-miR-4326 |
| hsa-miR-182-5p | hsa-miR-532-3p | hsa-miR-140-3p | hsa-miR-676-3p | hsa-miR-204-5p | hsa-miR-10a-5p | hsa-miR-3909 |
| hsa-miR-18a-3p | hsa-miR-5586-3p | hsa-miR-129-2-3p | hsa-miR-6761-3p | hsa-miR-214-5p | hsa-miR-211-5p | hsa-miR-4484 |
| hsa-miR-18b-3p | hsa-miR-615-5p | hsa-miR-146a-3p | hsa-miR-6806-3p | hsa-miR-145-5p | hsa-miR-216a-5p | hsa-miR-769-3p |
| hsa-miR-191-5p | hsa-miR-625-3p | hsa-miR-30c-1-3p | hsa-miR-6843-3p | hsa-miR-188-5p | hsa-miR-23b-5p | hsa-miR-625-3p |
| hsa-miR-193a-5p | hsa-miR-6505-3p | hsa-miR-296-3p | hsa-miR-26b-3p | hsa-miR-200c-5p | hsa-miR-125a-5p | hsa-miR-4732-5p |
| hsa-miR-23a-5p | hsa-miR-652-5p | hsa-miR-361-3p | hsa-miR-493-3p | hsa-miR-381-5p | hsa-miR-128-2-5p | hsa-miR-1287-5p |
| hsa-miR-25-5p | hsa-miR-660-5p | hsa-miR-365a-3p | hsa-miR-149-3p | hsa-miR-383-5p | hsa-miR-361-5p | |
| hsa-miR-3074-5p | hsa-miR-671-3p | hsa-miR-365b-3p | hsa-let-7d-3p | hsa-miR-330-5p | hsa-miR-362-5p | |
| hsa-miR-30c-1-3p | hsa-miR-676-5p | hsa-miR-331-3p | hsa-miR-217-3p | hsa-miR-584-5p | hsa-miR-365b-5p | |
| hsa-miR-3170- | hsa-miR-6875-3p | hsa-miR-18b-3p | hsa-miR-135a-2-3p | hsa-miR-616-5p | hsa-miR-328-5p | |
| hsa-miR-3183- | hsa-miR-6875-5p | hsa-miR-431-3p | hsa-miR-125b-2-3p | hsa-miR-642a-5p | hsa-miR-331-5p | |
| hsa-miR-320a-3p | hsa-miR-6892-3p | hsa-miR-500a-3p | hsa-miR-134-3p | hsa-miR-659-5p | hsa-miR-339-5p | |
| hsa-miR-330-3p | hsa-miR-708-5p | hsa-miR-501-3p | hsa-miR-136-3p | hsa-miR-1296-5p | hsa-miR-433-5p | |
| hsa-miR-339-3p | hsa-miR-760- | hsa-miR-502-3p | hsa-miR-150-3p | hsa-miR-939-5p | hsa-miR-502-5p | |
| hsa-miR-345-3p | hsa-miR-766-3p | hsa-miR-504-3p | hsa-miR-188-3p | hsa-miR-1180-5p | hsa-miR-652-5p | |
| hsa-miR-345-5p | hsa-miR-769-5p | hsa-miR-548b-3p | hsa-miR-151a-3p | hsa-miR-3677-5p | hsa-miR-874-5p | |
| hsa-miR-361-5p | hsa-miR-877-5p | hsa-miR-33b-3p | hsa-miR-433-3p | hsa-miR-3682-5p | hsa-miR-885-5p | |
| hsa-miR-937-5p | hsa-miR-92a-2-5p | hsa-miR-1296-3p | hsa-miR-193b-3p | hsa-miR-4661-5p | hsa-miR-216b-5p | |
| hsa-miR-99b-5p | | hsa-miR-1271-3p | hsa-miR-552-3p | hsa-miR-4709-5p | hsa-miR-1229-5p | |

**Supplementary Table S6** Prediction scores for 3D structures of influential miRNA-mRNA pairs in fusion positive KIRP cases. Color font is based onnetwork modules. Greay background represents downregulated genes and upregulated microRNAs.

**iPTM** =interface predicted template modelling; **PTM** =predicted template modelling

| mRNA | miRNA | iPTM score | PTM score |
|---|---|---|---|
| SOD2 | hsa-miR-874-3p | 0.84 | 0.9 |
| NQO1 | hsa-miR-130a-5p | 0.83 | 0.9 |
| SOD2 | hsa-miR-30c-2-3p | 0.81 | 0.92 |
| NQO1 | hsa-miR-30c-1-3p | 0.81 | 0.91 |
| NQO1 | hsa-miR-30c-2-3p | 0.72 | 0.91 |
| SOD2 | hsa-miR-30c-1-3p | 0.71 | 0.9 |
| G6PD | hsa-miR-30c-2-3p | 0.63 | 0.88 |
| SOD2 | hsa-miR-497-5p | 0.46 | 0.89 |
| GPX4 | hsa-miR-874-5p | 0.45 | 0.89 |
| SOD2 | hsa-miR-26b-3p | 0.37 | 0.87 |
| NQO1 | hsa-miR-874-5p | 0.36 | 0.85 |
| ATP6V1C1 | hsa-miR-874-5p | 0.9 | 0.89 |
| ATP6V1C1 | hsa-miR-874-3p | 0.87 | 0.92 |
| ATP6V1C1 | hsa-miR-195-5p | 0.86 | 0.9 |
| ATP6V1C1 | hsa-miR-26b-3p | 0.84 | 0.92 |
| FLCN | hsa-miR-27b-3p | 0.83 | 0.91 |
| FNIP2 | hsa-miR-497-5p | 0.82 | 0.9 |
| ATP6V1F | hsa-miR-30c-1-3p | 0.79 | 0.9 |
| ATP6V1F | hsa-miR-195-5p | 0.65 | 0.9 |
| FNIP2 | hsa-miR-130a-3p | 0.64 | 0.89 |
| FNIP2 | hsa-miR-200b-5p | 0.63 | 0.89 |
| ATP6V1C1 | hsa-miR-497-5p | 0.54 | 0.88 |
| FNIP2 | hsa-miR-130a-5p | 0.52 | 0.88 |
| STAT1 | hsa-miR-874-5p | 0.85 | 0.91 |
| STAT1 | hsa-miR-874-3p | 0.84 | 0.91 |
| STAT1 | hsa-miR-10a-5p | 0.84 | 0.92 |
| STAT1 | hsa-miR-27b-5p | 0.77 | 0.9 |
| LAMP1 | hsa-miR-130a-5p | 0.57 | 0.89 |
| CCND1 | hsa-miR-185-3p | 0.84 | 0.91 |
| CCND1 | hsa-miR-130b-3p | 0.82 | 0.91 |
| MAPK10 | hsa-miR-185-5p | 0.81 | 0.91 |
| MAPK11 | hsa-miR-148a-5p | 0.77 | 0.9 |
| MAPK10 | hsa-miR-185-3p | 0.7 | 0.91 |
| MAPK10 | hsa-miR-342-5p | 0.69 | 0.9 |
| MAPK10 | hsa-miR-130b-5p | 0.61 | 0.9 |
| MAPK10 | hsa-miR-148a-5p | 0.49 | 0.89 |
| CCND1 | hsa-miR-148a-5p | 0.46 | 0.88 |
| MAPK10 | hsa-miR-148a-3p | 0.39 | 0.87 |
| MAPK10 | hsa-miR-221-3p | 0.35 | 0.87 |
| F11R | hsa-miR-342-5p | 0.9 | 0.92 |
| MARVELD2 | hsa-miR-342-5p | 0.89 | 0.92 |
| F11R | hsa-miR-130b-5p | 0.85 | 0.9 |
| OCLN | hsa-miR-221-3p | 0.83 | 0.9 |
| F11R | hsa-miR-185-5p | 0.81 | 0.91 |
| MARVELD2 | hsa-miR-148a-3p | 0.73 | 0.89 |
| OCLN | hsa-miR-185-3p | 0.6 | 0.89 |
| MARVELD2 | hsa-miR-185-5p | 0.49 | 0.88 |
| MARVELD2 | hsa-miR-130b-3p | 0.49 | 0.87 |
| MARVELD2 | hsa.miR-5589-5p | 0.47 | 0.89 |
| F11R | hsa-miR-5589-5p | 0.32 | 0.86 |

**Supplementary Table S7.** Prediction scores for 3D structures of influential miRNA-mRNA pairs in fusion negative KIRP cases

| miRNA | mRNA | iPTM score | PTM score |
|---|---|---|---|
| BRCA1 | hsa-mir-660-3p | 0.88 | 0.92 |
| BRCA1 | hsa-mir-135a-2-3p | 0.84 | 0.92 |
| BRCA1 | hsa-mir-483-3p | 0.64 | 0.89 |
| BRCA1 | hsa-mir-654-5p | 0.52 | 0.88 |
| BRCA1 | hsa-mir-370-3p | 0.29 | 0.86 |
| BUB1B | hsa-mir-483-5p | 0.52 | 0.87 |
| CCNB2 | hsa-mir-483-5p | 0.87 | 0.91 |
| CCNB2 | hsa-mir-654-5p | 0.8 | 0.91 |
| CDH1 | hsa-mir-3127-5p | 0.75 | 0.9 |
| CDH1 | hsa-mir-25-5p | 0.63 | 0.89 |
| CDH1 | hsa-mir-6892-3p | 0.55 | 0.88 |
| CXCL8 | hsa-mir-370-3p | 0.38 | 0.84 |
| ESR1 | hsa-mir-6892-5p | 0.85 | 0.91 |
| ESR1 | hsa-mir-3127-5p | 0.75 | 0.92 |
| ESR1 | hsa-mir-6892-3p | 0.73 | 0.91 |
| ESR1 | hsa-mir-885-3p | 0.44 | 0.89 |
| FOS | hsa-mir-1295a | 0.84 | 0.91 |
| RAD51 | hsa-mir-483-3p | 0.73 | 0.9 |
| RAD51 | hsa-mir-135a-2-3p | 0.42 | 0.88 |
| RAD51 | hsa-mir-370-3p | 0.4 | 0.85 |
| TLR2 | hsa-mir-660-3p | 0.89 | 0.92 |
| TLR2 | hsa-mir-483-5p | 0.73 | 0.88 |
| VEGFA | hsa-mir-6509-5p | 0.7 | 0.9 |
| VEGFA | hsa-mir-1293 | 0.55 | 0.89 |
| VEGFA | hsa-mir-25-5p | 0.42 | 0.87 |

**iPTM** =interface predicted template modelling; **PTM** =predicted template modelling
